# Structure-Activity Relationship Study Identifies a Novel Lipophilic Amiloride Derivative that Efficiently Kills Chemoresistant Breast Cancer Cells

**DOI:** 10.1101/2023.05.25.542364

**Authors:** Michelle Hu, Ruiwu Liu, Noemi Castro, Liliana Loza Sanchez, Julie Learn, Ruiqi Huang, Kit S. Lam, Kermit L. Carraway

**Author notes:** Corresponding author: Kermit L. Carraway UC Davis School of Medicine 4645 2^nd^ Avenue, room 1100B Sacramento, CA 95817.

## Abstract

Derivatives of the potassium-sparing diuretic amiloride are preferentially cytotoxic toward tumor cells relative to normal cells, and have the capacity to target tumor cell populations resistant to currently employed therapeutic agents. However, a major barrier to clinical translation of the amilorides is their modest cytotoxic potency, with estimated IC_50_ values in the high micromolar range. Here we report the synthesis of ten novel amiloride derivatives and the characterization of their cytotoxic potency toward MCF7 (ER/PR-positive), SKBR3 (HER2-positive) and MDA-MB-231 (triple negative) cell line models of breast cancer. Comparisons of derivative structure with cytotoxic potency toward these cell lines underscore the importance of an intact guanidine group, and uncover a strong link between drug-induced cytotoxicity and drug lipophilicity. We demonstrate that our most potent derivative called LLC1 is preferentially cytotoxic toward mouse mammary tumor over normal epithelial organoids, acts in the single digit micromolar range on breast cancer cell line models representing all major subtypes, acts on cell lines that exhibit both transient and sustained resistance to chemotherapeutic agents, but exhibits limited anti-tumor effects in a mouse model of metastatic breast cancer. Nonetheless, our observations offer a roadmap for the future optimization of amiloride-based compounds with preferential cytotoxicity toward breast tumor cells.

## Introduction

While amiloride has been used clinically for decades as an anti-kaliuretic in the management of hypertension [1], and in the lab as a pharmacological tool for examining sodium transport [2,3], substantial *in vitro* and *in vivo* data point to its anti-cancer and anti-metastatic potential [4]. High concentrations of amiloride have been demonstrated to inhibit mutagen-induced carcinogenesis [5–7], tumor formation [8–10] and metastatic progression [11,12] in rats and mice. These outcomes are generally ascribed to the cytostatic effects of amiloride inhibition of sodium hydrogen exchanger-1 (NHE1), and to the motility-suppressing effects of urokinase plasminogen activator (uPA) inhibition [5].

Beyond cytostatic and anti-invasiveness activities, more recent observations indicate that amiloride and its derivatives also exhibit tumor cell-selective cytotoxicity via a non-apoptotic mechanism [13–16]. For example, our studies with the derivative 5-(N,N-hexamethylene)amiloride (HMA) underscore the notion that the properties of amiloride derivatives may be ideally suited for targeting particularly aggressive or therapy-refractory tumors [17,18]. HMA efficiently kills bulk breast tumor cells independent of tumor subtype, proliferation state or species of origin, but does not efficiently kill non-transformed cells derived from a variety of tissues at the same concentrations. Indeed, cell lines derived from diverse tumor types are equally susceptible to HMA, suggesting that its mechanism of cytotoxic action may be dependent on cellular transformation rather than patient-specific genetic alterations [17,18]. Moreover, HMA is cytotoxic toward breast cancer cell populations that are resistant to currently-employed therapies, including the highly refractory cancer stem cell population. Finally, HMA induces morphological changes within the lysosomes of tumor cells, and cytotoxicity is rescued by an inhibitor of the lysosomal protease cathepsin [17,18], pointing a central role for the lysosome in its mechanism of action.

While our previous *in vitro* studies have revealed many attractive properties of amiloride derivatives, and suggest that derivatization of the C(5) position of its pyrazine ring can markedly enhance the tumor-selective cytotoxic properties of the drug, further translation of HMA is hampered by its relatively modest potency [17,18] and poor pharmacokinetic properties in mice [19]. In this study we aimed to glean further insight into structure-activity relationships (SARs) within the amiloride pharmacophore by comparing the cytotoxic potential of derivatives modified at the C(2) and C(5) positions. At the same time, we sought to identify more potent derivatives that optimize tumor cell-specific cytotoxicity and preserve the ability to target chemoresistant populations. Here we report the synthesis and characterization of 10 novel amiloride derivatives, and identify a novel lead compound that acts with significantly greater potency than HMA that preserves tumor-selective cytotoxicity and efficiently eradicates chemoresistant breast tumor cells.

## Results

### Lipophilic additions to the amiloride pharmacophore increase cytotoxic potency

Amiloride is often modified at the C(2), C(5), and C(6) positions of its pyrazine ring (see Fig. 1A) to improve its transporter inhibition activity and selectivity [8,20], and such derivatives have been employed to assess the contribution of ion channel function to tumor growth and progression [4,14–19,21–29]. However, our recent studies indicating that the amiloride derivative HMA can provoke the non-apoptotic death of tumor cells relative to non-cancer cells *in vitro* by acting via the lysosome [17,18] prompt the question of whether these cytotoxic properties might be further optimized.

**Figure 1.**
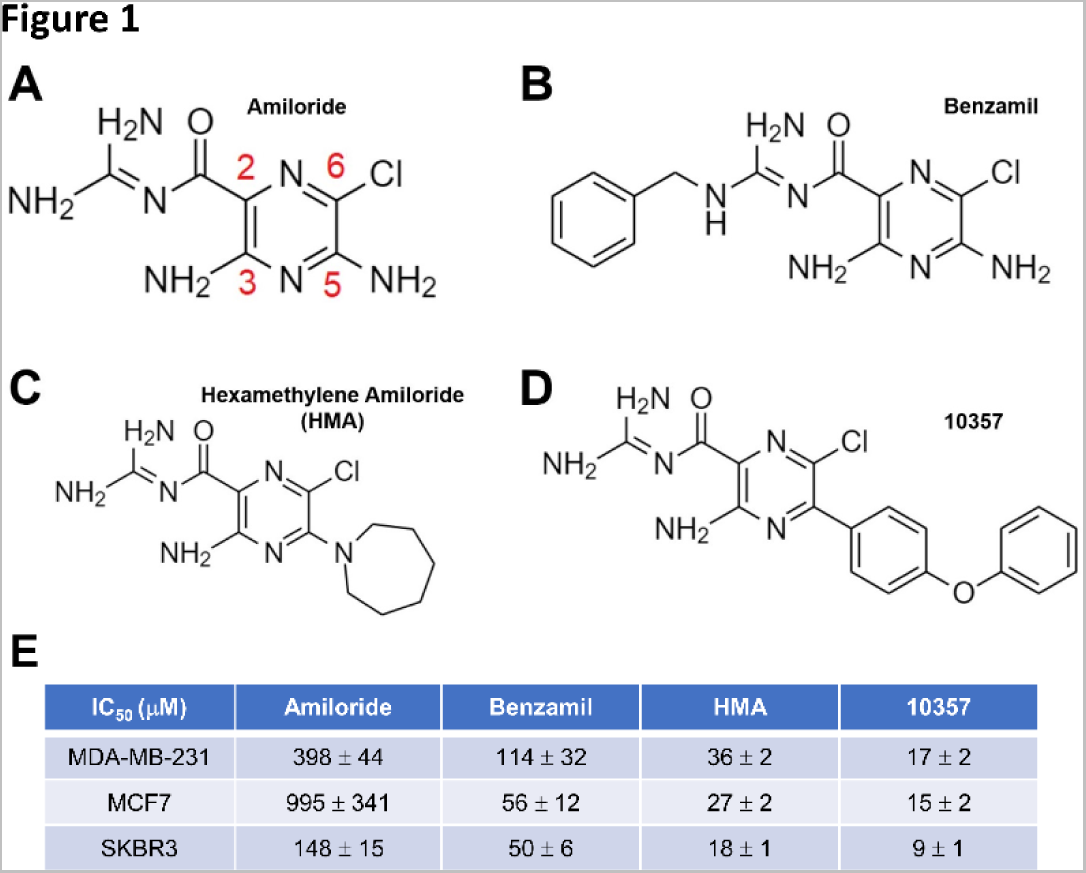
Modification of amiloride with lipophilic substituents enhances its cytotoxic potency. **(A)** Structure of amiloride with its pyrazine carbons numbered. **(B-D)** Structures of 2- and 5-substituted amiloride derivatives benzamil, HMA and 10357 are depicted. **(E)** Cytotoxic potency of amilorides toward cultured breast cancer cell lines after 24-hour treatment. Data are presented as IC_50_ (μM) averages ± SEM.

As HMA adds a lipophilic hexamethylene group to the core pharmacophore at the C(5) position, we first asked whether lipophilic modification increases the cytotoxic potency of amiloride using previously published amiloride derivatives. Fig. 1 depicts the structures of parent drug amiloride (Fig. 1A), C(2) derivative benzamil (Fig. 1B), C(5) derivative HMA (Fig. 1C), and C(5) derivative 10357 (Fig. 1D) [26], and summarizes the cytotoxic potency of these derivatives toward cell lines representing ER/PR-positive (MCF7), HER2-positive (SKBR3) and triple-negative (MDA-MB-231) breast tumor subtypes in an MTT cell viability assay (Fig. 1E). Consistent with previous observations [14,15,17,28,30,31], we observed that while IC_50_ values for amiloride cytotoxicity are in the hundreds of micromolar, the three previously published lipophilic derivatives are 3- to 66-fold more potent, and followed the general efficacy pattern 10357 > HMA > benzamil > amiloride toward each of the cell line models. These observations are consistent with the notion that membrane permeability and access to an intracellular target(s) may be key to the cytotoxic mechanism of action.

### Novel amiloride derivatives underscore the importance of logP and the guanidine group

To more rigorously test the link between amiloride derivative lipophilicity and cytotoxic potential, we synthesized nine novel amiloride derivatives, most modified with various substituents at the C(5) position. The structures and IC_50_ values for cytotoxicity toward the three breast cancer cell line models are summarized in Table 1, and the chemical characteristics of all derivatives ordered by estimated IC_50_ are summarized in Table 2. Three key points emerge from comparisons of these derivatives. First, Table 2 points to a strong correlation between cytotoxic efficacy and lipophilicity, reflected in partition coefficient (logP) values of the derivatives. For example, LLC1 and 10357, which exhibit some of the highest logP values (2.46 and 2.88, respectively) among the group, also exhibit the most potent cytotoxicity toward the breast cancer cell lines (5-13 µM and 9-17 µM, respectively). On the other hand, the low logP values for amiloride and LLC10 (-0.89 and -0.28, respectively) correlate with particularly poor cytotoxicity (148-995 µM and 883-2732 µM, respectively). Spearman rank correlation coefficients of IC_50_ with logP were 0.0083, 0.0035, and <0.0001, respectively for MDA-MB-231, MCF7 and SKBR3 cells. Second, a direct comparison of derivatives LLC1 and LLC3 (Table 1) illustrates the importance of maintaining the integrity of the C(2) guanidine structure. Both derivatives contain an identical lipophilic C(5) substituent, but the removal of one of the guanidine amines results in a 3- to 23-fold decrease in cytotoxic potency. Finally, to a first approximation each of the derivatives acts with similar potency toward all three cell lines (Table 1). This suggests that, consistent with previous observations [17,18], amiloride derivatives act with similar efficacy toward the different breast cancer subtypes. Overall, these observations reveal new parameters for the optimization of amiloride derivatives, and highlight LLC1 as a novel lead.

**Table 1.**
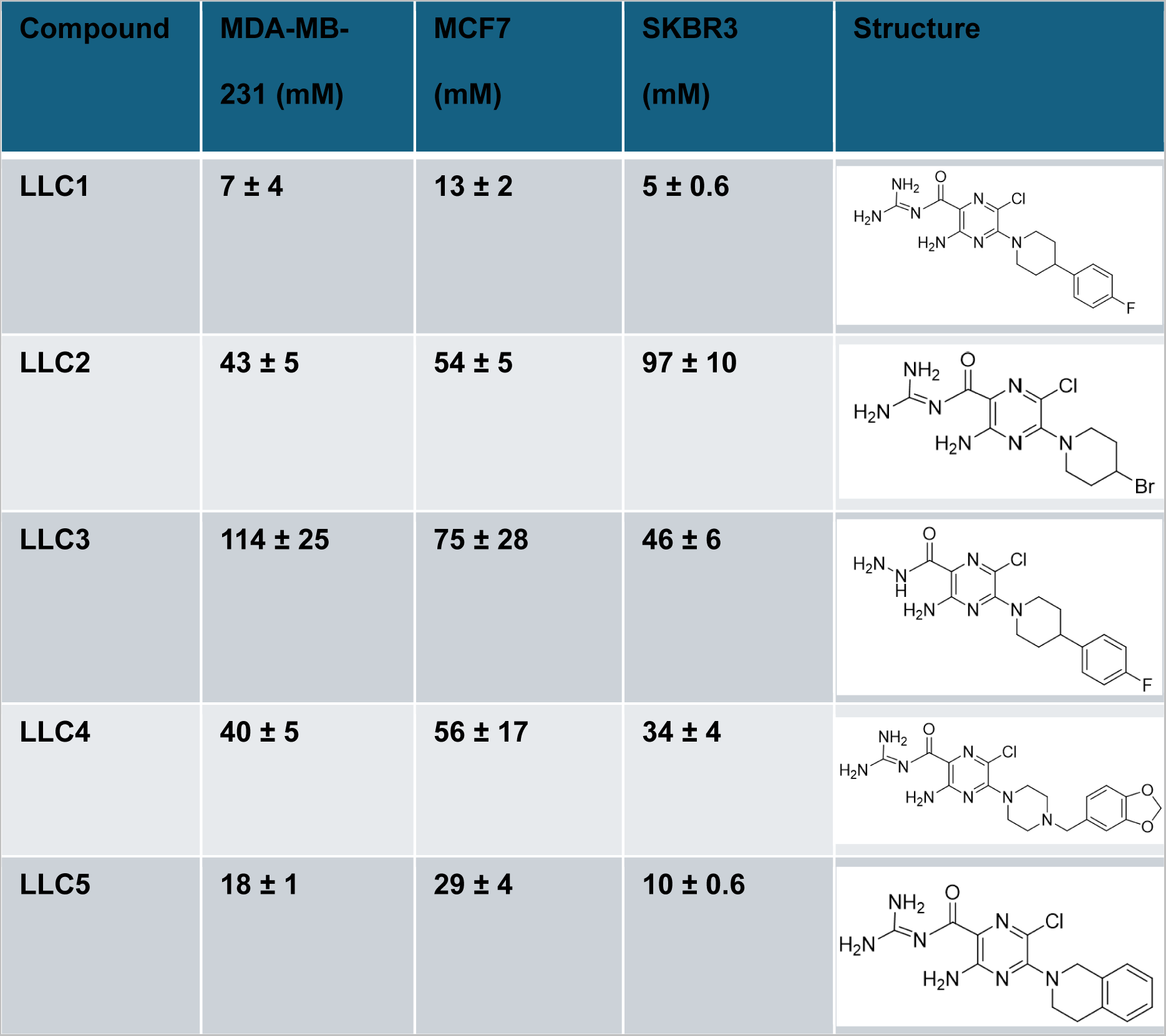
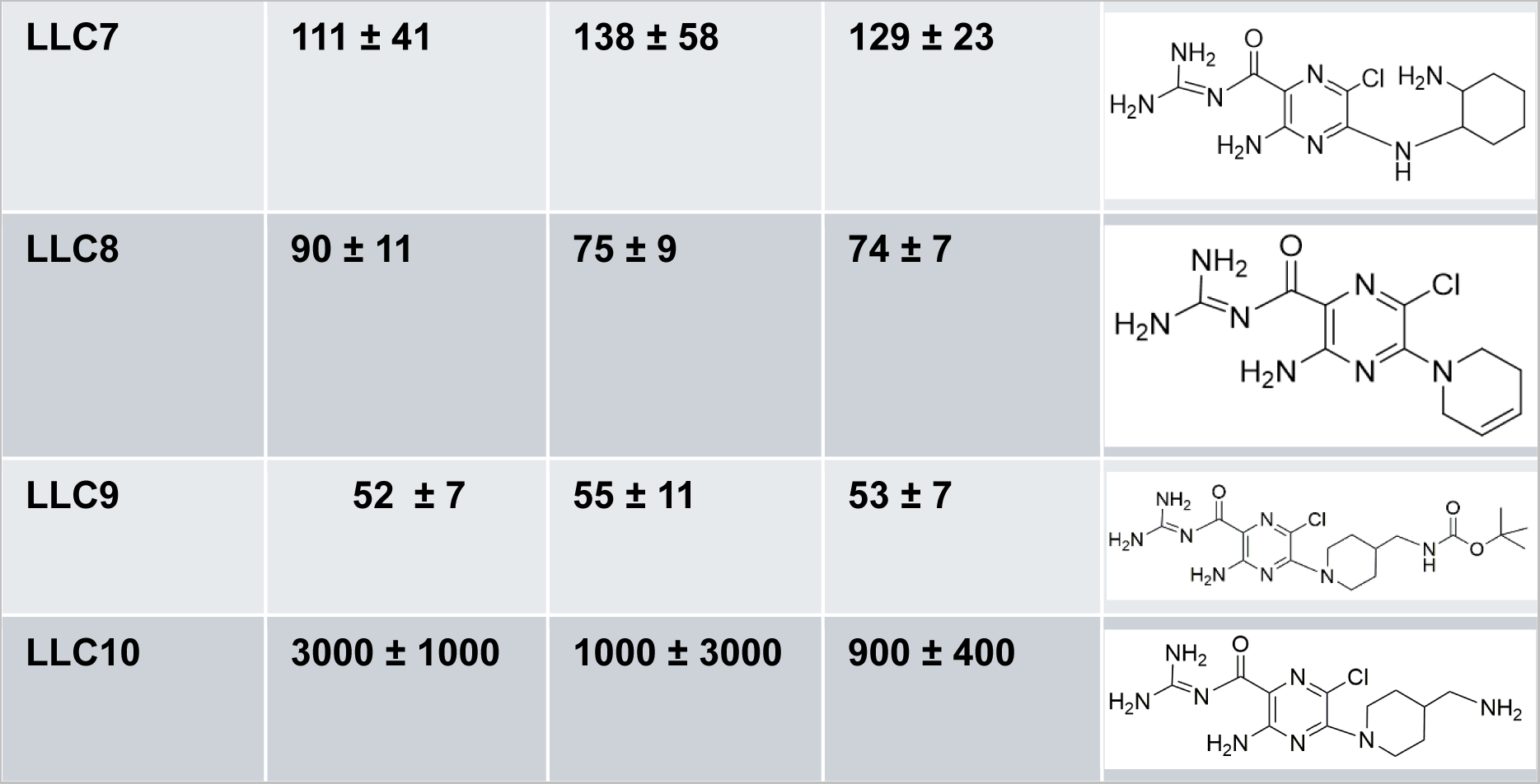
Potencies of novel 2- & 5-substituted amiloride derivatives. Calculated IC_50_ data from MDA-MB-231, MCF7, and SKBR3 cells (24-hour treatment), and specific structures for each compound are represented in the table. Data are compiled from at least three biological replicates of each cell line and are presented as averages ± SEM.

| Compound | MDA-MB-231 (mM) | MCF7 (mM) | SKBR3 (mM) | Structure |
| --- | --- | --- | --- | --- |
| LLC1 | $7 \pm 4$ | $13 \pm 2$ | $5 \pm 0.6$ | |
| LLC2 | $43 \pm 5$ | $54 \pm 5$ | $97 \pm 10$ | |
| LLC3 | $114 \pm 25$ | $75 \pm 28$ | $46 \pm 6$ | |
| LLC4 | $40 \pm 5$ | $56 \pm 17$ | $34 \pm 4$ | |
| LLC5 | $18 \pm 1$ | $29 \pm 4$ | $10 \pm 0.6$ | |
| LLC7 | 111 ± 41 | 138 ± 58 | 129 ± 23 |  |
| LLC8 | 90 ± 11 | 75 ± 9 | 74 ± 7 |  |
| LLC9 | 52 ± 7 | 55 ± 11 | 53 ± 7 |  |
| LLC10 | 3000 ± 1000 | 1000 ± 3000 | 900 ± 400 |  |

**Table 2.**
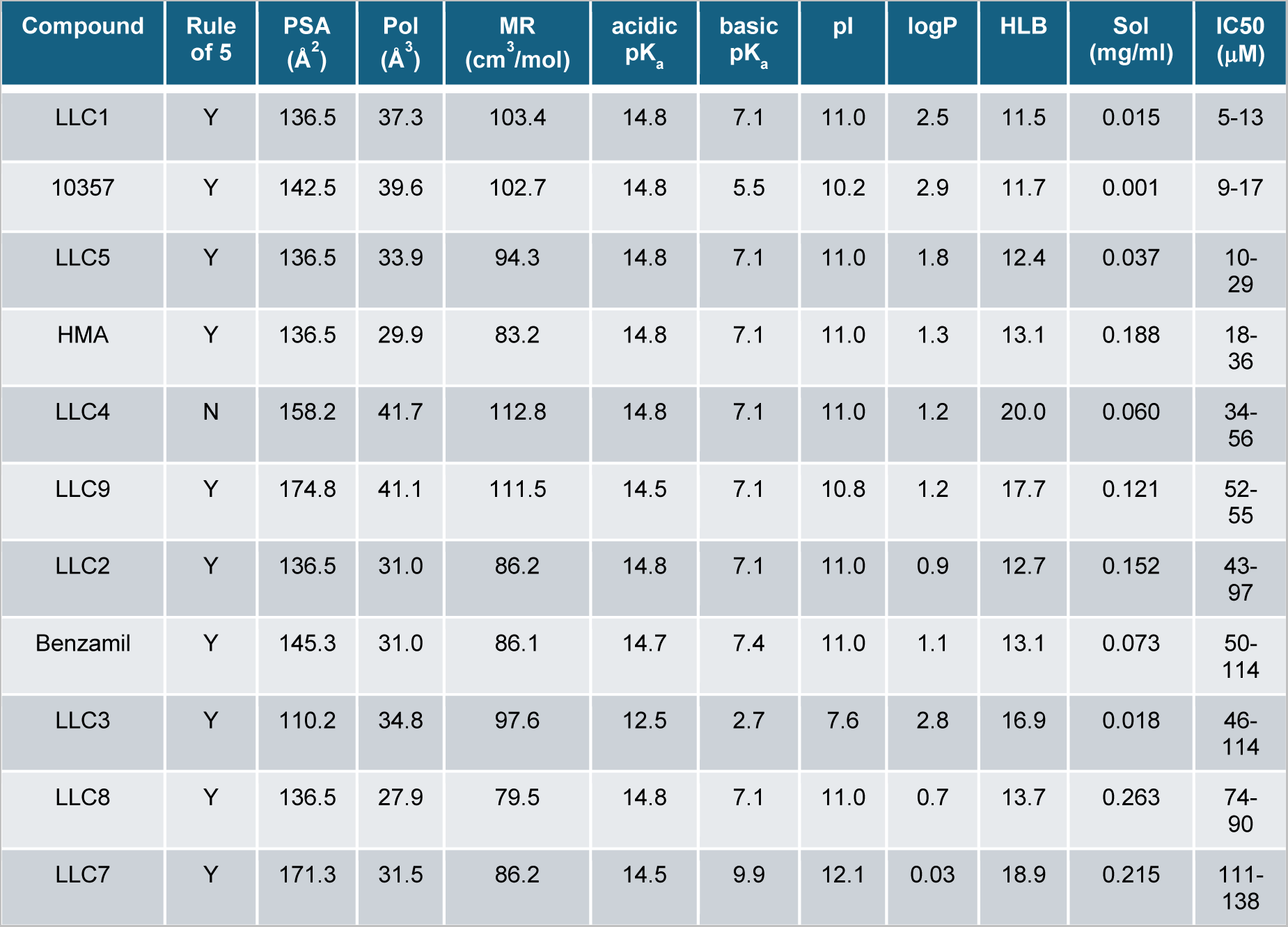
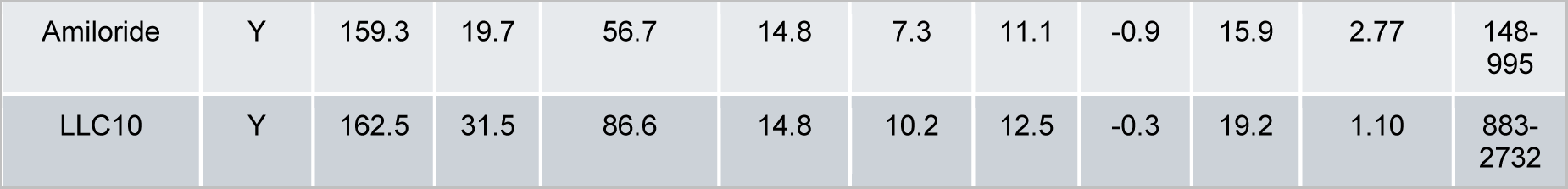
Chemical and cytotoxicity characteristics of amiloride derivatives. The chemical properties of the novel amiloride derivatives are ordered according to collective cytotoxicity IC_50_ toward MCF7, SKBR3 and MDA-MB-231 breast cancer cell lines (far right column). Chemical data [Rule of 5, topological polar surface area (PSA), polarizability (Pol), molar refractivity (MR), acidic pKa, basic pKa, isoelectric point (pI), logP, hydrophilic-lipophilic balance (HLB), and intrinsic solubility (Sol)], were obtained using Chemicalize (https://chemicalize.com, developed by ChemAxon).

### LLC1 preferentially reduces the viability of murine mammary tumor organoids ex vivo

We have previously demonstrated that HMA reduces the viability of organoids derived from the PymT and NDL genetically-engineered mouse models of breast cancer [18]. To determine whether LLC1 is selectively cytotoxic toward mammary tumors relative to normal tissue, we compared its impact on tumor and normal mammary gland organoids generated from Balb/cJ mice orthotopically engrafted with 4T1 (ER^-^/PR^-^/HER2^-^), an isogenic murine mammary cancer cell line. The approximate IC_50_ value of LLC1 toward human tumor cell lines (10 µM, Table 1) was used to compare the efficiency of cell death as a function of time. We observed that while LLC1 reduces the viability of tumor organoids over 72 hours of treatment (Figs. 2A and 2B), normal mammary gland organoids tend to be resistant to LLC1 treatment (Figs. 2C and 2D). These observations suggest that, similar to its predecessor HMA [17,18], LLC1 exhibits significant selectivity toward cancer cells and leaves untransformed cells largely unharmed.

**Figure 2.**
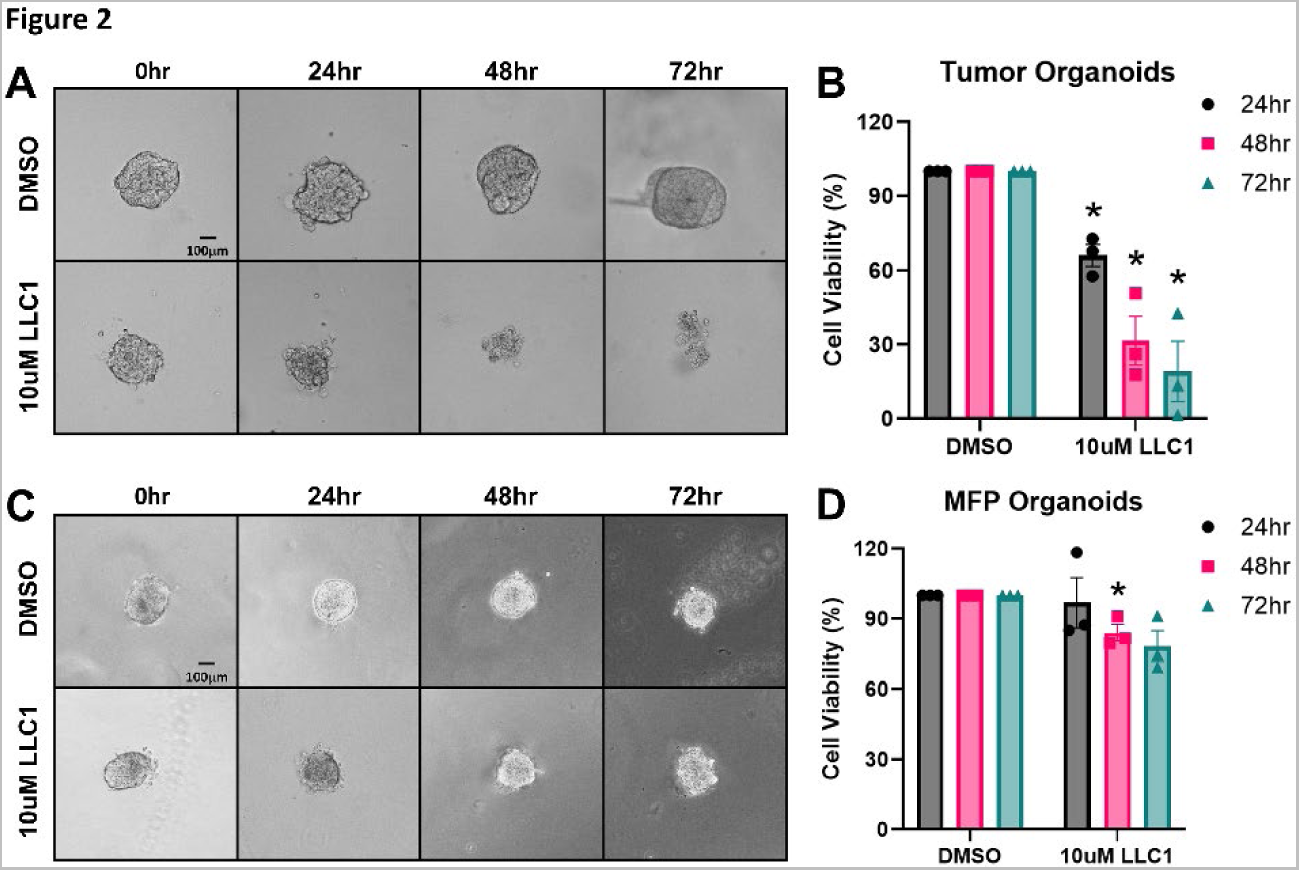
LLC1 selectively reduces the viability of mouse mammary tumors *ex vivo*. Representative images of LLC1-treated (10 µM) organoids generated from mammary tumors **(A)** and normal mammary glands **(C)** from 4T1-engrafted Balb/cJ mice are shown. Scale bar = 100 µm. **(B, D)** Organoids were treated with 10 µM LLC1, and viability was assessed using RealTime Glo over 72 hours and normalized to vehicle treatment (DMSO). Tumor **(B)** or mammary fat pad (MFP) **(D)** organoid viability was monitored every 24 hours. Data were compiled from three biological replicates of organoids derived from tumors or mammary glands of independent mice. Data are presented as averages ± SEM and significance assessed by t-test. *, *P* < 0.05.

### LLC1 acts on chemoresistant breast cancer cell populations

Our previous observations indicate that HMA is cytotoxic toward ‘persister’ populations of breast cancer cells that survive 3-9 day exposure to conventional chemotherapeutic agents cisplatin, docetaxel and doxorubicin (DOXO) [18]. To confirm HMA’s capabilities and test LLC1’s properties, we examined the impact of both drugs on the viability of various breast cancer cell line models of short-term and established therapeutic resistance. We first examined the response of chemoresistant populations of MDA-MB-231 cells toward the derivatives. In the experiment shown in Fig. 3, cells were first pretreated for 48 hours with DMSO vehicle, and then further treated for 24 hours with DMSO, 250 nM docetaxel (DTX), 5 μM DOXO, and either 40 μM HMA (Fig. 3A) or 10 μM LLC1 (Fig. 3D). In each case, drug treatment lowered cell viability by 15-40%, confirming that the drugs behave as expected. However, in cells pretreated for 48 hours with either DTX (Figs. 3B and 3E) or DOXO (Figs. 3C and 3F), a second challenge with the pretreating drug did not reduce viability while HMA or LLC1 treatment lowered viability by 25-60%, consistent with the interpretation that chemotherapy-resistant populations are sensitive to amiloride derivatives.

**Figure 3.**
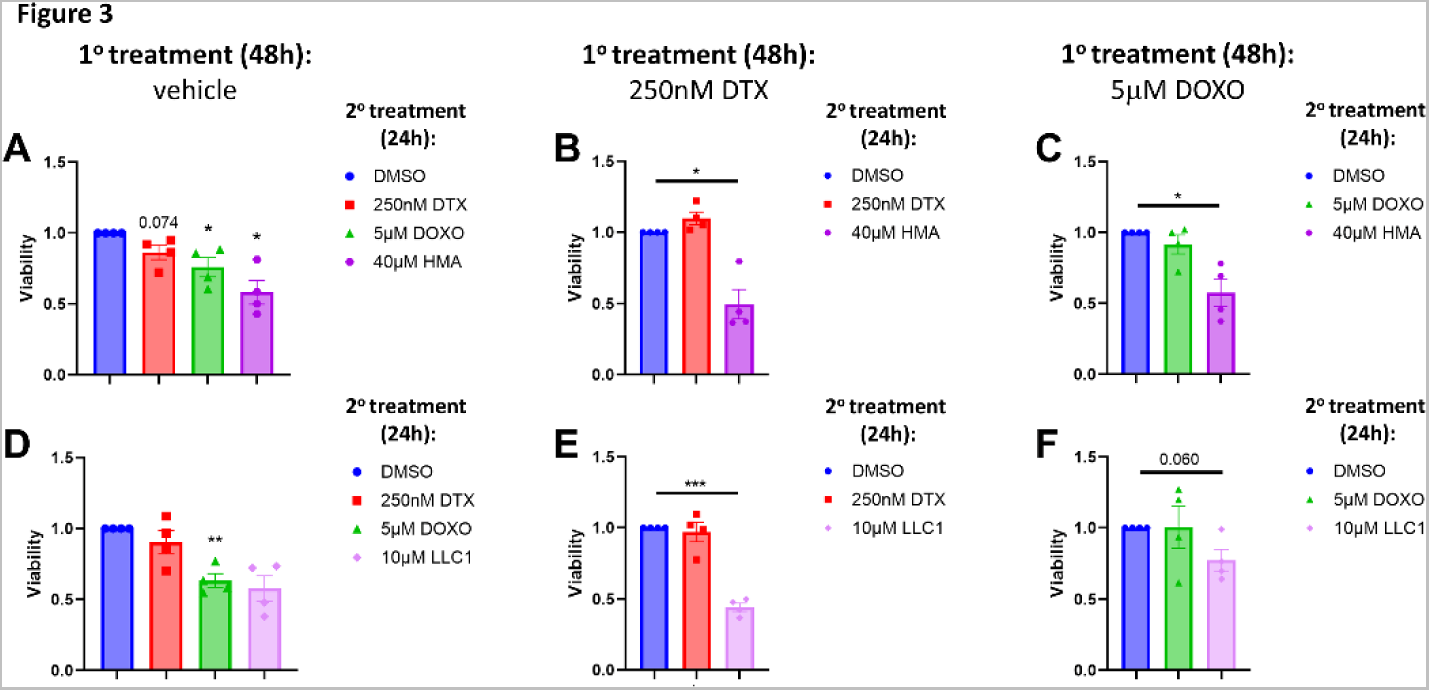
HMA and LLC1 are cytotoxic toward chemoresistant triple-negative breast cancer cell populations. **(A,D)** MDA-MB-231 cells were initially treated with vehicle (DMSO) for 48 hours, followed by a second challenge with DMSO, 250 nM docetaxel (DTX), 5 µM doxorubicin (DOXO), or either 40 µM HMA **(A)** or 10 µM LLC1 **(D)** for 24 hours, and then cell viability was determined by MTT assay. **(B,C,E,F)** Following 48 hours 250 nM DTX **(B,E)** or 5 µM DOXO **(C,F)** pretreatment, residual cells were treated with vehicle, initial chemotherapy treatment (DTX or DOXO), HMA **(B,C)**, or LLC1 **(E,F)** for an additional 24 hours. Data represent a minimum of four biological replicates and are presented as averages ± SEM, and significance was assessed by t-test. *, *P* < 0.05; **, *P* < 0.01; ***, *P* < 0.001.

We further observed that roughly half of T47D cells, an ER/PR^+^ breast cancer cell line, exhibit resistance to cell death induced by 72-hour treatment with doxorubicin (DOXO, Fig. 4A) and docetaxel (Fig. 4D), reflected in the inability of high doses of either drug to further impact viability in a titration. However, we found that both HMA and LLC1 are cytotoxic toward the residual population that remains after chemotherapeutic treatment (Figs. 4B, 4C and 4E, 4F), and exhibit similar IC_50_ values as cell lines representing a variety of solid tumor types (Table 1).

**Figure 4.**
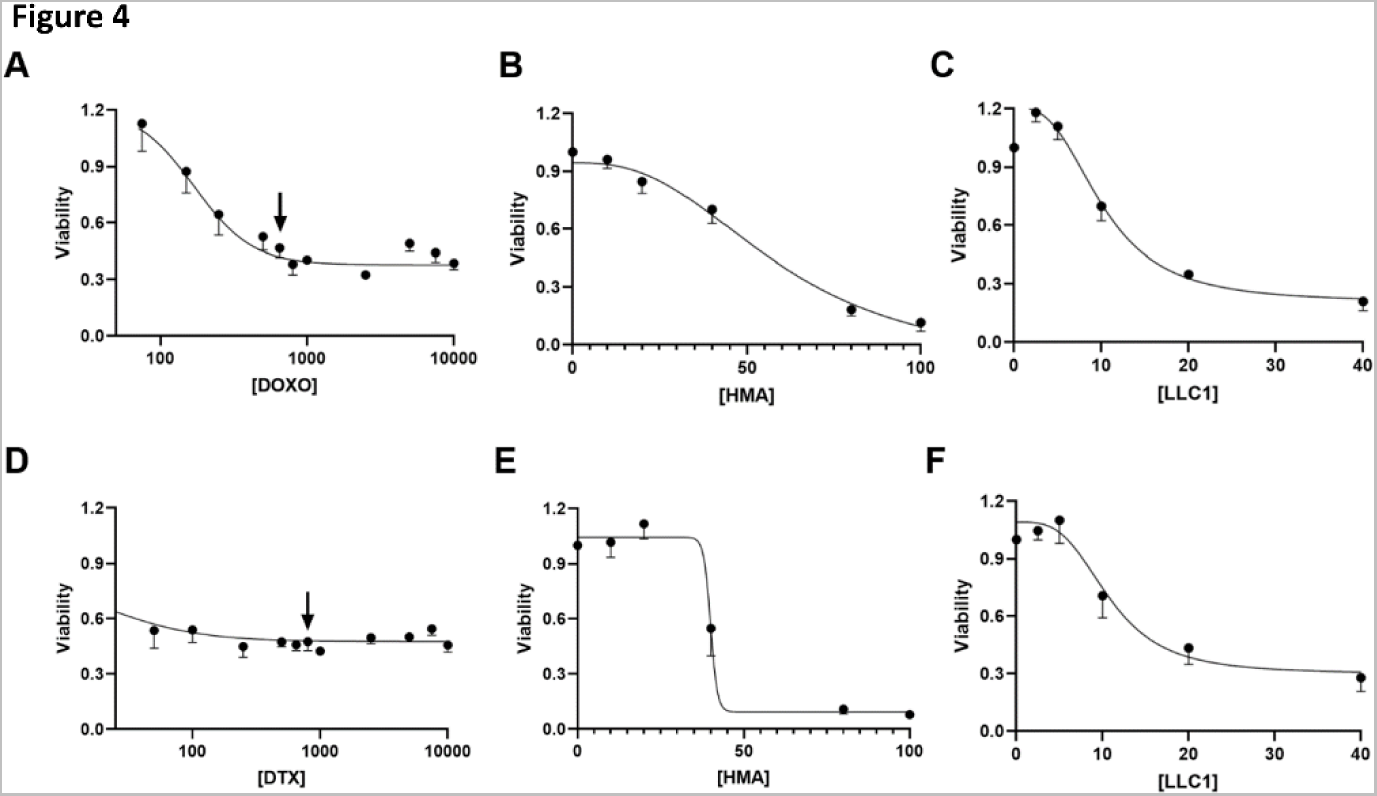
HMA and LLC1 are cytotoxic toward chemoresistant ER/PR-positive breast cancer cell populations. Dose-response curves of T47D cells treated with doxorubicin (DOXO) **(A)** or docetaxel (DTX) **(D)** for 72 hours are depicted. Populations of cells that persist at drug concentrations higher than 650 nM DOXO and 800 nM DTX (arrows) are considered drug resistant. Residual T47D cells that survive treatment with these levels of drugs for 72 hours were then titrated with HMA **(B,E)** or LLC1 **(C,F)** for an additional 24 hours and viability assessed. Data are compiled from at least three biological replicates.

Additionally, we observed that a breast cancer cell line selected for stable therapeutic resistance is as sensitive to amiloride derivatives as the drug-sensitive parental line. MX-100 [32] is a derivative of MCF7 cells selected for stable mitoxantrone resistance that overexpresses ABCG2 (aka breast cancer resistance pump, BCRP) by 30-fold. MCF7 TR-1 and MCF7 TR-5 [33] are stable tamoxifen resistant derivatives, while MCF TS is (z)-4-hydroxytamoxifen (TAM) sensitive. Each cell line was treated with varying concentrations of TAM, DTX, DOXO, HMA, and LLC1, and cytotoxicity IC_50_ values were determined (Table 3). We observed that parental MCF7 cells are 3-13 fold more sensitive to chemotherapeutic agents DTX and DOXO than MX-100, but both cell lines are similarly sensitive to HMA and LLC1. Likewise, MCF7 TS cells are far more sensitive to tamoxifen than MCF7 TR-1 and MCF7 TR-5 cells, but all three lines are similarly sensitive to HMA and LLC1. Collectively, these observations provide strong evidence that amiloride derivatives act on breast tumor cell populations that are resistant to clinically pertinent anti-cancer agents.

**Table 3.** MCF7 chemoresistant cells are susceptible to amiloride derivatives but not conventional chemotherapeutics. The table depicts parent and chemoresistant MCF7 cell line derivatives titrated with chemotherapeutic agents tamoxifen (TAM), docetaxel (DTX), doxorubicin (DOXO), HMA, or LLC1 for 24 hours to determine their IC_50_ values. ND denotes that an IC_50_ value could not be determined due to the very high dosage required. Data represent a minimum of three independent biological replicates and are presented as averages ± SEM.

| Cell Line | TAM (mM) | DTX (mM) | DOXO (mM) | HMA (mM) | LLC1 (mM) |
| --- | --- | --- | --- | --- | --- |
| MCF7 | - | 1 ± 0.6 | 2 ± 0.3 | 27 ± 2 | 13 ± 2 |
| MCF7 MX-100 | - | 13 ± 3 | 7 ± 0.5 | 39 ± 3 | 12 ± 1 |
| MCF7 TS | 13 ± 1 | - | - | 68 ± 3 | 25 ± 1 |
| MCF7 TR-1 | ND | - | - | 67 ± 4 | 26 ± 2 |
| MCF7 TR-5 | ND | - | - | 59 ± 5 | 19 ± 2 |

Finally, we observed that both HMA and LLC1 are as cytotoxic toward diverse mammalian cell line models of cancer as human breast cancer cell line models. In the experiments summarized in Supplementary Table S3, we assessed the cytotoxic potencies of HMA and LLC1 toward Met-1 (triple-negative), NDL (HER2-positive), and 4T1 (triple-negative) murine breast cancer cells, as well as UCDK9MM3 (melanoma), UCDK9OSA29 (osteosarcoma), and D17 (osteosarcoma) canine tumor cell lines. We observed that HMA provokes the death of all models with IC_50_ values in the 20-40 μM range, while LLC1 induces cytotoxicity in the 7-14 μM range. These values are consistent with their potencies toward human breast cancer cell lines (Table 1), collectively providing support for the notion that amiloride derivatives act to eradicate transformed cells in a tumor type-, subtype-, and species-agnostic manner.

### LLC1 provokes little observable tissue toxicity or anti-tumor effects in vivo

As a more potent cytotoxic agent than HMA, we sought to assess the *in vivo* anti-tumor efficacy of LLC1 using the orthotopic, immune-intact 4T1-Balb/c model of metastatic triple-negative breast cancer. We began by performing a maximum tolerated dose (MTD) study with Balb/cJ mice, where animals were intraperitoneally injected with 0 mg/kg, 15 mg/kg, 30 mg/kg, and 45 mg/kg LLC1 three times over the course of one week. In this experiment we observed no obvious fluctuations in body weight (Fig. 5A) or aberrant behaviors in any of the animals. Likewise, bloodwork revealed no significant changes at any LLC1 treatment level in markers of liver and kidney damage compared to control (0 mg/kg LLC1) or the reference values (Fig. 5B). Moreover, organs including spleen, heart, liver, and kidneys exhibited no gross morphological (Figs. 5C-F) or histopathological differences (Supplementary Fig. S15) with LLC1, suggesting that under the employed treatment conditions LLC1 is well tolerated by mice.

**Figure 5.**
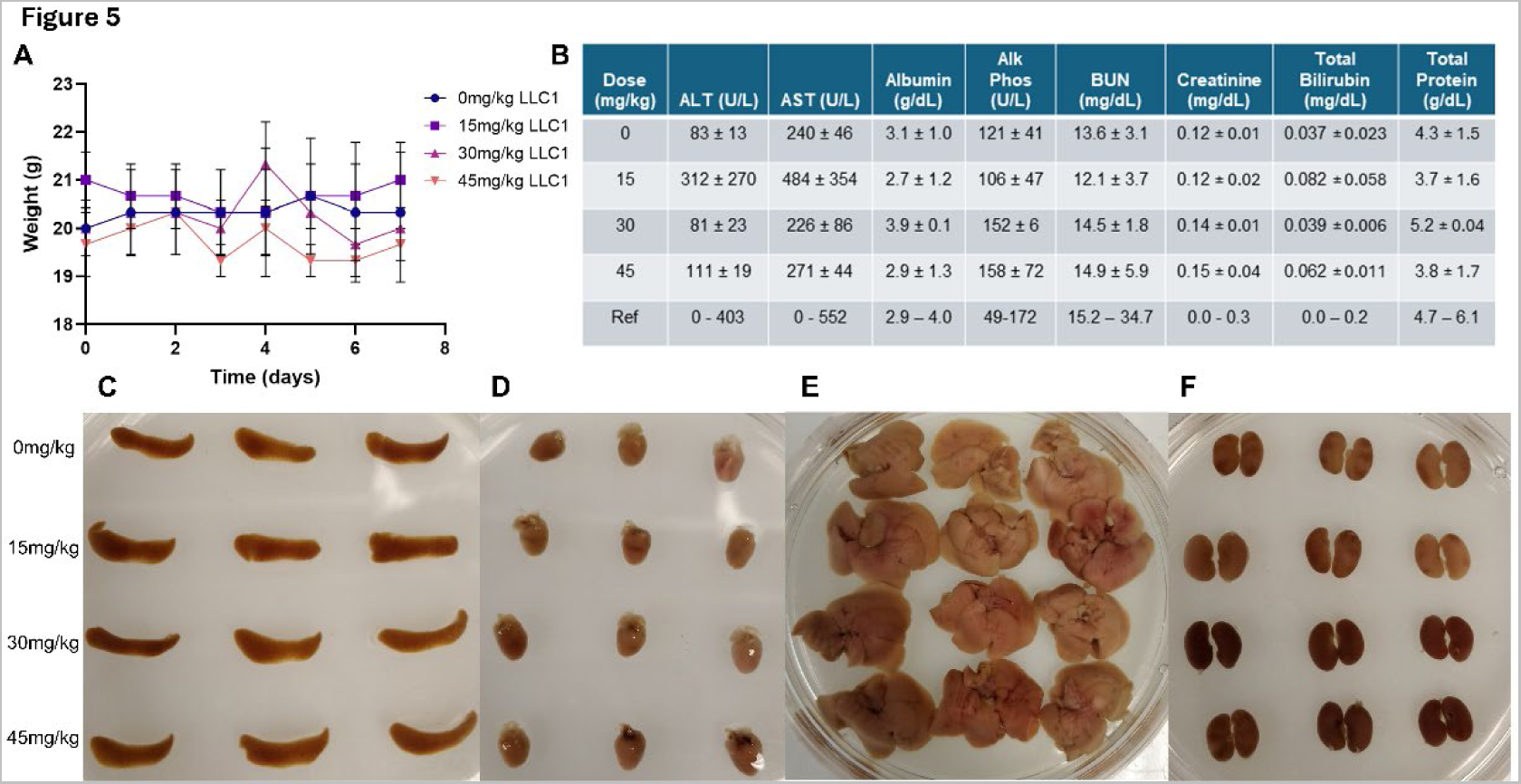
LLC1 is well tolerated by mice. Adult Balb/cJ mice were injected intraperitoneally with 0-45 mg/kg LLC1 on days 0, 3 and 6 of the study and sacrificed on day 7. **(A)** Body weights were recorded throughout the study. **(B)** A blood chemistry panel illustrates the effects of repeated LLC1 treatment on liver (ALT, alanine transaminase; AST, aspartate transaminase; albumin; alk phos, alkaline phosphatase; total bilirubin; total protein) and kidney (BUN, blood urea nitrogen; creatinine; total protein) markers. Images of mouse **(C)** spleen, **(D)** heart, **(E)** liver, and **(F)** kidneys are displayed for each dose after formalin fixation.

To assess the impact of LLC1 on tumor growth properties we orthotopically implanted 4T1 cells into the mammary fat pads of Balb/cJ mice. A treatment paradigm of intraperitoneally-delivered vehicle or 30mg/kg LLC1 three times per week for three weeks was initiated after tumors reached 100mm^3^ in volume (Fig. 6A), and animals were sacrificed a day after the last dose. Tumor volume (Fig. 6B) and mouse body weight (Supplementary Fig. S16) were monitored over the course of treatment, and tumor volume (Fig. 6C), number of metastatic lesions observed per lung lobe (Fig. 6D), and the area of metastatic coverage (Fig. 6E), were recorded at the experimental endpoint. We observed no statistically significant differences in tumor growth and metastatic characteristics upon treatment of animals with LLC1. Importantly, inspection of primary tumors revealed no evidence of increased necrotic cell death with LLC1 treatment, strongly suggesting that drug dosing or delivery was insufficient to impact tumor characteristics.

**Figure 6.**
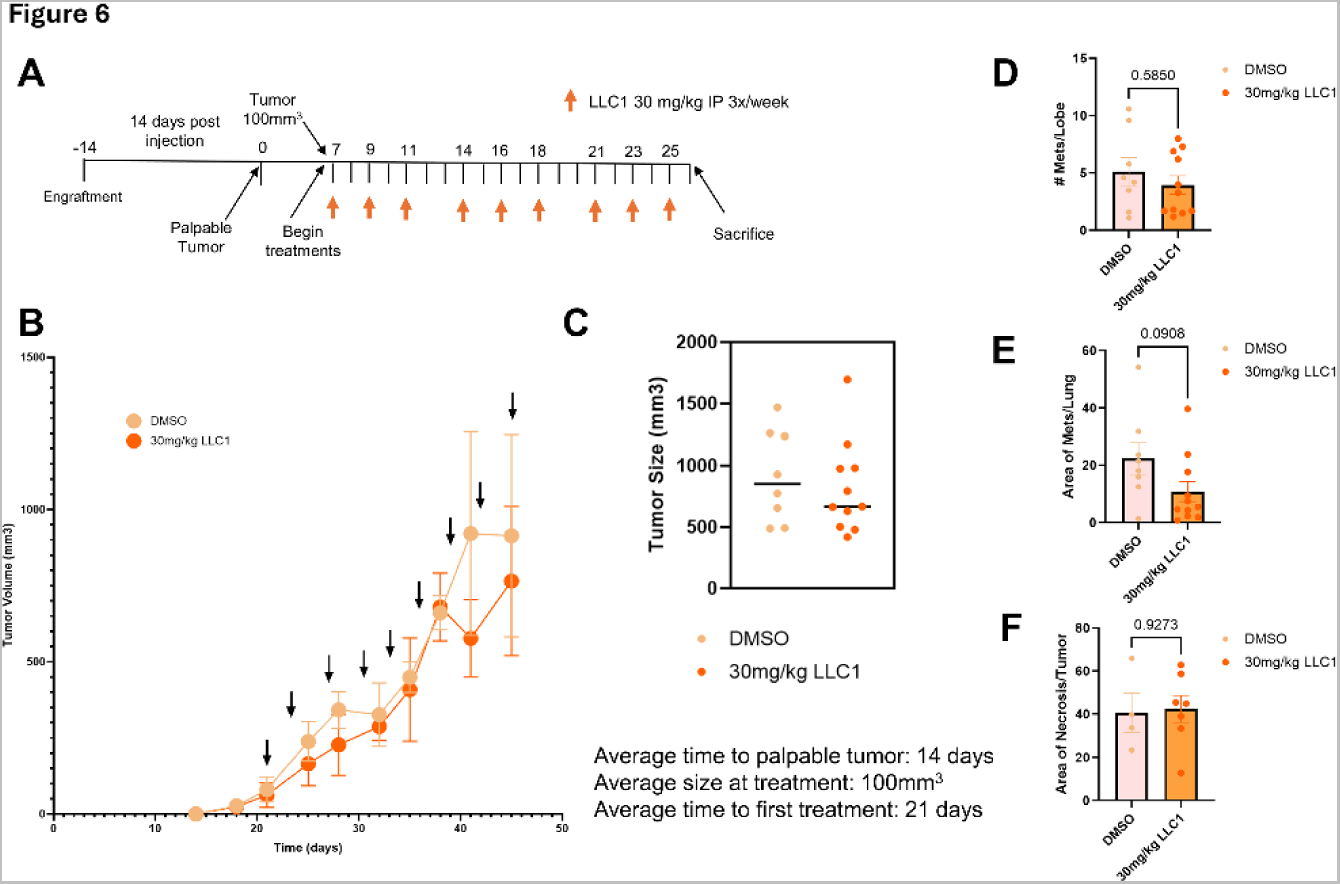
LLC1 exhibits limited therapeutic efficacy toward a 4T1-Balb/cJ model of metastatic breast cancer. **(A)** Schematic of the treatment regimen is illustrated, where animals are treated three times per week for three weeks with vehicle (n=8) or 30mg/kg LLC1 (n=11) after tumors reach 100mm^3^ in volume, and sacrificed 19 days after the initial injection. **(B)** Tumor volumes were determined for animals treated with vehicle and LLC1. Averages ± SEM at each time point are depicted. **(C)** Tumor volumes at sacrifice are compared. Metastatic burden was established through analysis of metastatic lesions per lobe **(D)** and metastatic area of the lung **(E)**. **(F)** Area of necrosis was evaluated between vehicle- and LLC1-treated cohorts. Differences in metastatic burden parameters and necrotic area were compared by t-test.

## Discussion

Amiloride derivatives offer significant promise as anticancer therapeutic agents because they exhibit tumor cell-selective cytotoxicity and harbor the potential to eradicate chemoresistant tumor cell populations [18]. However, modest potency and poor pharmacokinetic properties have hampered efforts to translate these agents. The optimization of amiloride structure is an enticing area for medicinal chemists and biologists alike because the relative ease of synthesis and chemical modification allow the straightforward development of new derivatives while optimizing its unique anticancer properties. Our study reveals that LLC1, a novel lipophilic C(5) amiloride derivative, induces cell death in the single digit micromolar range, selectively kills tumor organoids relative to normal mammary gland organoids, and eradicates chemoresistant breast cancer cell populations. Collectively, our observations demonstrate that systematic analysis of amiloride pharmacophore modification can lead to improved anticancer drug candidates, agents that have the capacity to act across many tumor types, subtypes and even species.

Importantly, our SAR analysis provides two key insights into building more effective cancer-selective cytotoxic amiloride derivatives. First, we observed a strong relationship between amiloride lipophilicity, reflected in more positive logP values, and drug cytotoxicity. Indeed, our most potent derivative LLC1 exhibits one of the highest logP values of the compounds we have examined, suggesting that further increases in lipophilicity could result in even more effective drug leads. These conclusions dovetail with current trends in drug development that prioritize more lipophilic drugs because of their higher likelihood of passing safety and ADME concerns during Phase 1 clinical studies [34]. Second, disruption of the guanidine moiety at the C(2) position appears deleterious to cytotoxic function. LLC1 and LLC3 are identical except for the deletion of one of the guanidine amino groups in LLC3, and this change leads to an order of magnitude loss of potency. In this regard it might be interesting to determine whether lipophilic derivatives of other existing guanidine-containing drugs, or lipophilic compounds containing multiple guanidine groups, might similarly exhibit tumor cell-selective cytotoxicity and improved efficacy.

Our findings strongly suggest that amilorides act as cationic amphiphilic drugs (CADs), a broad class of agents defined by the presence of both a lipophilic moiety and a hydrophilic ionizable amine [16,35,36]. Over five dozen FDA-approved agents fit the general structural definition of CADs, including antidepressants, antibiotics, antiarrhythmics, and diuretics [16,36]. CADs characteristically accumulate in lysosomes and frequently provoke lysosome-mediated lipidosis which can be followed by cell death, and for this reason investigators will often disregard hits with these agents in drug screens [35]. However, the tumor-selective cytotoxicity of this drug class, together with observations that lipidotic responses may be readily reversed upon CAD withdrawal [35,36], underscore the existence of a therapeutic window that might be exploited for cancer patient benefit.

Their high degree of hydrophobicity allows CADs to spontaneously cross membranes, while the low pH of the lysosome facilitates drug protonation, luminal trapping and accumulation, and disruption of lysosomal hydrolase activities. Enzymes of the lysosomal sphingolipid catabolic cascade appear particularly susceptible to CADs, possibly because electrostatic interactions of these enzymes and their chaperones with highly acidic intraluminal vesicles required for enzyme function are shielded by the positively charged agents [35,36]. The resulting accumulation of sphingolipid substrates then destabilizes the lysosomal limiting membrane, leading to lysosomal membrane permeabilization (LMP) and lysosome-dependent cell death (LDCD) [37–40]. A variety of cellular stressors, including cationic amphiphilic drugs, microtubule inhibitors, nanoparticles and lysosomotropic detergents, can damage and permeabilize the limiting lysosomal membrane. Beyond compromising lysosome function, LMP leads to the release of lysosomal hydrolases that act on cytosolic substrates to elicit a cascade of events culminating in cytolysis, the defining event in necrotic cell death [40]. Importantly, sequestration of CADs at the site of their targets in the lysosome and shielded from drug resistance-mediating plasma membrane drug pumps whose expression is common to many chemoresistant tumor cells may contribute to the sensitivity of chemoresistant cells to amiloride derivatives.

Two features of LMP/LDCD make the engagement of this cell death mode particularly attractive for anti-cancer therapeutics development. First, lysosomal membranes of transformed cells are inherently more fragile than those of normal cells [38,41], explaining tumor cell selectivity of amiloride derivatives and offering a potential therapeutic window that might be exploited clinically. Second, we have observed both previously and in this study that chemoresistant cells, including cancer stem cells (CSCs), are as susceptible to this form of cell death as are differentiated (bulk) cancer cells [40]. CSCs are the rare subpopulation of tumor cells responsible for both tumor recurrence and metastasis, are notoriously resistant to chemotherapeutic and targeted therapeutic agents [42,43], and remain a vexing barrier to achieving substantially improved patient outcomes. The similar susceptibility of CSCs and bulk cancer cells to LMP/LDCD suggests that CSC subpopulations cannot easily escape LMP/LDCD-inducing therapeutic agents to initiate primary or metastatic recurrence through reversion to the differentiated state. This then raises the possibility that these agents could be significantly more effective in altering patient outcomes than existing anti-cancer drug classes.

The limited anti-tumor effects of LLC1 *in vivo* is disappointing but not surprising. Previous studies demonstrate that HMA has a very short half-life in mice [19], and in our previous studies we were only able to observe significant anti-tumor effects of HMA when we encapsulated the drug in a disulfide-linked nanoparticle that markedly prolongs half-life [18]. Since we strongly suspect that drug elimination is a central issue with LLC1, future studies will be aimed at identifying higher potency amiloride derivatives with more favorable pharmacokinetic properties.

## Materials and Methods

### Cell Culture and Drug Treatments

Human breast cancer cell lines MDA-MB-231, MCF7, SKBR3, and T47D, and mouse mammary cancer cell line 4T1, were purchased from American Type Culture Collection (Manassas, VA, USA) and maintained at 37°C and 10% CO_2_ in Dulbecco’s Modified Eagle Medium (DMEM) supplemented with 10% fetal bovine serum (Genesee Scientific) and antibiotics (penicillin/streptomycin; Gibco - Thermo Fisher). MCF7-B7-TS (MCF7 TS), MCF7-G11-TR-1 (MCF7 TR-1), MCF7-G11-TR-5 (MCF7 TR-5), and MCF7 MX-100 cells were obtained from the Physical Sciences-Oncology Network Bioresource Core Facility at ATCC or gifted from Dr. A.M. Yu, UC Davis. The tamoxifen (TAM) resistant lines were maintained in MCF7 base media supplemented with 1 μM or 5 μM TAM [44]. The mitoxantrone resistant line (MX-100) was maintained as previously described [45]. Met-1 (gifted by A.D. Borowsky, UC Davis) and NDL murine cells were maintained as previously described [46,47]. UCDK9MM3, UCDK9OSA27, and D17 canine cells (gifted by R.B. Rebhun, UC Davis) were maintained as previously described [48–50]. Cell line attributes are summarized in Supplementary Table S1.

Amiloride, HMA, benzamylamiloride (benzamil), and amiloride derivative 10357 were sourced as described in Supplementary Table S2, applied to cells at 50-70% confluency in DMSO (final vehicle concentration <0.5%), and analysis of amiloride derivative cytotoxicity was carried out after 24 hours except in the case of the organoid experiment (up to 72 hours). In some experiments cells were titrated (Fig. 4) with chemotherapeutic agents (docetaxel, 0-10μM; doxorubicin, 0-10μM), or were pretreated with agents (docetaxel, 0.65 μM; doxorubicin, 0.8 μM) to pre-select for short-term drug resistance prior to treatment with amiloride derivatives (Fig. 3).

### Design and Synthesis of Novel Amiloride Derivatives

The new amiloride derivatives were designed to introduce novel hydrophobic groups at the C(5) position to improve cell permeability and potential *in vivo* availability. Synthesis of LLC1 was carried out as shown in Supplementary Fig. S1. To a suspension of methyl 3-amino-5,6-dichloro-2-pyrazinecarboxylate (compound **1**; 444 mg, 2.0 mmol) in anhydrous dimethylformamide (DMF; 3 mL) was added 4-(4-fluorophenyl)piperidine (compound **2**; 358.5 mg, 2.0 mmol) and *N*,*N*-Diisopropylethylamine (697 µL, 4.0 mmol), the reaction mixture was stirred at room temperature overnight and then added to cold water. The precipitate was collected by centrifuge and washed with water. The crude product (compound **3**) was used for the next step without further purification. 0.5 M NaOCH_3_ solution in CH_3_OH (20 mL) was added to guanidine hydrochloride (955.3 mg, 10 mmol) and the resulting mixture was stirred at room temperature for 0.5 hr. The white precipitate was removed by filtration, and the free guanidine solution in methanol was added to a suspension of methyl ester (compound 3; 364.8mg, 1.0 mmol) in DMF (5 mL) and stirred at room temperature overnight. Brine (20 mL) was added, and the mixture extracted with ethyl acetate (3 × 30 mL). The organic layer was washed with 10% NaCl (2 × 30 mL), dried over anhydrous Na_2_SO_4_, and concentrated under reduced pressure. The crude mixture was dissolved in 5 mL of 30% CH_3_CN/70% H_2_O and 0.1% trifluoroacetic acid and purified by a reverse phase HPLC. The collected eluent was lyophilized to yield the designed product LLC1. The structure was verified by Orbitrap high resolution ESI-MS: calculated 392.1402, found 392.1391 [M+1] (Supplementary Fig. S2A). The purity was >98% as shown in HPLC (Supplementary Fig. S2B). LLC3 was synthesized by mixing compound **3** in DMF with 1.0 M hydrazine (10 eq.) in ethanol overnight (Supplementary Fig. S1). The chemical identity and purity were verified by ESI-MS and HPLC (Supplementary Figs. S3A and 3B), respectively.

Other LLC compounds were synthesized using a similar approach with modification. Different amines were used to displace the C5-Cl in compound **1**: 4-bromopiperidine hydrobromide (**4**) for LLC2 and LLC8, -piperonylpiperazine (**6**) for LLC4, 1,2,3,4-tetrahydroisoquinoline (**7**) for LLC5, 1,2-diaminocyclohexane (**8**), tert-butyl methyl(piperidin-4-ylmethyl)-carbamate (**12**) for LLC9 and LLC10. The synthetic approach of LLC2 and LLC8 is shown in Fig. S4. LLC4, LLC5 and LLC7 were prepared as shown in Fig. S5. The synthesis of LLC9 and LLC10 is shown in Fig. S6. Treatment of LLC9 with 50% trifluoroacetic acid in dichloromethane (DCM) for 1 hour yielded LLC10. All LLC compounds have purity >95% as shown in HPLC and their identities were verified by HR ESI-MS (Figs. S7-S13).

### MTT Assay

Cells were seeded on 24-well plates and cultured to 50-70% confluency prior to treatment with various compounds for 24-72 hour depending on the assay context. Following treatments, media was aspirated from cells and replaced with 5 mg/mL 3-(4,5-Dimethyl-2-thiazolyl)-2,5-diphenyl-2H-tetrazolium bromide (MTT, #M5655, Sigma Aldrich) solution in base media. Cells were incubated at 37°C for 1-2 hours to induce formazan crystal formation, then washed with phosphate-buffered saline (PBS) to remove impurities. Crystals were dissolved using isopropyl alcohol (0.5% 1N HCl in isopropanol), and sample absorbance (λ_ex_ 570 nm) was measured with a FilterMax F5 microplate reader (Molecular Devices) and Multi-Mode Analysis software (Version 3.4.0.27, Beckman Coulter). The reported IC_50_ values for each drug were calculated from the concentration required to induce half-maximal suppression of absorbance across 3-4 biological replicates.

### Ex Vivo Drug Cytotoxicity Organoid Assay

Tumors were harvested after 1 x 10^6^ 4T1 cells engrafted into Balb/cJ mice (#000651, The Jackson Laboratory) at the 4^th^ mammary fat pad were grown to 1.0-1.5 cm in diameter. Tumors or pooled mammary glands from Balb/cJ mice were chopped into fine pieces using a razor blade before addition to digestion buffer (DMEM/F12 #SH30023, HyClone;1% PenStrep, Gibco - Thermo Fisher; 2% fetal bovine serum (FBS), Genesee Scientific; collagenase/hyaluronidase (5%: tumors & 10%: mammary glands), #07912, StemCell Technologies; 5 mg/mL insulin, Gibco - Thermo Fisher) and vortexed. Samples were digested for 2hr at 37°C with gentle shaking, and samples vortexed every 30 minutes, before they were processed. After neutralization of the digestion buffer with DMEM (Gibco - Thermo Fisher) containing 10% FBS, organoids were centrifuged at 700 rpm to remove cellular debris. ACK lysis buffer (Gibco - Thermo Fisher) was used to remove red blood cells from the organoids. A quick spin was performed to remove additional debris and to pellet organoids. Once resuspended in PBS, organoids were embedded into MatriGel (#354230, Corning) and cultured in organoid growth media (OGM) on 24-well plates for 24 hours as described previously [51]. After overnight incubation in OGM, organoids were treated with vehicle or 10 µM LLC1 for up to 72 hours, and viability was measured using RealTime Glo (#G9711, Promega) according to manufacturer’s instructions. Images were taken before drug treatment (0 hour) and every 24 hour over the course of drug treatment. Representative brightfield images were taken with an Olympus IX81 microscope with CellSens Entry software. Chemiluminescent images were taken with a ChemiDoc Touch Imaging System (BioRad) and analyzed with FIJI (https://imagej.net/Fiji) software to quantify RealTime Glo signal.

### Statistical, Data, and Image Analysis

A minimum of three biological replicates per experiment was performed with values calculated and expressed as averages ± standard error of the mean (SEM), unless otherwise stated. Statistical significance was between control and test arms was determined using paired t-test; P-values of less than 0.05 were considered statistically significant. Data analysis was performed using Microsoft Excel or GraphPad Prism version 9.3.1 for Windows (GraphPad Software; San Diego, CA, USA; www.graphpad.com). Images were compiled in Microsoft PowerPoint and only brightness and contrast were altered for clarity.

### Maximum Tolerated Dose Study

All animal studies were performed in accordance with protocols approved by the Institutional Animal Care and Use Committee (IACUC) of the University of California, Davis. 8-10 week old Balb/cJ mice (000651) were purchased from The Jackson Laboratory and were acclimated to the animal room for at least one week prior to use. To determine the maximum tolerated dose (MTD) for LLC1, mice (n=3/cohort) were randomly separated into four groups and injected intraperitoneally with 0mg/kg (10% DMSO in saline), 15mg/kg, 30mg/kg, and 45mg/kg LLC1 in 10% DMSO in saline three times over seven days. Body weights and clinical signs were observed daily. Mice were sacrificed the day following the third treatment after cardiac puncture to collect blood. After coagulation and separation by centrifugation, serum was collected for blood chemistry panel analysis. Organs were harvested and fixed in 10% neutral buffered formalin for paraffin embedding and sectioning.

### LLC1 Therapeutic Efficacy Study

2x10^5^ 4T1 cells in PBS were mixed in a 1:1 volume/volume ratio with PuraMatrix Peptide Hydrogel (#354250, Corning) and injected bilaterally into the 4^th^ mammary fat pads of 8-10 week old Balb/cJ mice under anesthesia by continuous inhalation of 2% isoflurane gas. The 4^th^ nipple was used as a landmark for injection into the mammary gland with sterile tweezers lifting the nipple while the syringe needle loaded the cell suspension into the mammary fat pad. Tumor dimensions were measured twice per week with digital calipers and tumor volume calculated according to the formula v = (w^2^ x l) x 0.5236. Once tumors reached 100mm^3^, animals were randomized into two cohorts: 8 mice injected with 10% DMSO in saline and 11 mice injected with 30g/kg LLC1 in 10% DMSO in saline. Mice were injected intraperitoneally with their appropriate dose three times per week for three weeks before sacrifice by CO_2_ asphyxiation the day after the last treatment. Tumors and organs were harvested and fixed in 10% neutral buffered formalin for paraffin embedding and sectioning.

### Histology

H&E-stained sections of tumors, liver, kidney and heart were prepared as previously described [52]. 5µm sections were imaged using Keyence microscope BZ-X800, and LLC1 toxicity was assessed based on any morphologic changes. For the therapeutic study, all xenograft tumors were subjected to histological analysis. For tumors, the area of active necrosis was quantified and normalized to total tumor area from 2-6 randomly selected fields across three serial sections. Staining of lungs for metastatic lesions was modified from whole mount mammary gland and tumor protocols previously described [52]. Briefly, lungs were dehydrated, cleared, and stained with Mayer’s hematoxylin before imaging of macrometastases with a dissecting scope (Zeiss Stemi 2000-C; Axiocam ERc/5s). Analyses of lesions were performed using two approaches: 1) lesions were counted per lung lobe and averaged between two different slides, and 2) lesion area was quantified and averaged between total lung area. Lungs were then rehydrated and processed to confirm histology.

## Acknowledgements

We thank Dr. Frederic Gorin for providing compound 10357 and Dr. Aiming Yu for providing the MX-100 MCF7 derivative cell line. This research was funded by NIH/NCI grants R01CA250211, R01CA250211-S1, R01CA250211-S2, and R01CA230742-S1 (KLC). The UC Davis Combinatorial Chemistry and Chemical Biology Shared Resource is supported by NIH/NCI Cancer Center Support Grant P30CA093373 and was responsible for the synthesis and purification of the compounds. A preprint version of this article may be found at bioRxiv: https://www.biorxiv.org/content/10.1101/2023.05.25.542364v1.

## Author Contributions

Participated in research design: Hu, Liu, Carraway

Conducted experiments: Hu, Liu, Castro, Huang, Loza-Sanchez, Learn

Contributed new reagents or analytic tools: Liu, Lam

Performed data analysis: Hu, Liu

Wrote or contributed to the writing of the manuscript: Hu, Carraway

## Data Availability Statement

Data related to the synthesis and characterization of novel amiloride derivatives may be found in the Supplementary Materials. Other primary data presented in this study are available on request from the corresponding author.

