## Supplementary materials for "Structure-Activity Relationship Study Identifies a Novel Lipophilic Amiloride Derivative that Efficiently Kills Chemoresistant Breast Cancer Cells"

| Cell Line | Species | Subtype | ER | PR | HER2 | Tissue type | Source |
|---|---|---|---|---|---|---|---|
| MDA-MB-231 | Human | Basal | - | - | - | Metastatic Adenocarcinoma | ATCC #HTB-26 |
| MCF7 | Human | Luminal | + | + | - | Metastatic Adenocarcinoma | ATCC #HTB-22 |
| T47D | Human | Luminal | + | + | - | Invasive Ductal Carcinoma | ATCC #HTB-133 |
| SKBR3 | Human | Luminal | - | - | + | Adenocarcinoma | ATCC #HTB-30 |
| MCF7 TS | Human | Luminal | + | + | - | Metastatic Adenocarcinoma | Gifted by ATCC PBCF |
| MCF7 TR-1 | Human | Luminal | + | + | - | Metastatic Adenocarcinoma | Gifted by ATCC PBCF |
| MCF7 TR-5 | Human | Luminal | + | + | - | Metastatic Adenocarcinoma | Gifted by ATCC PBCF |
| MCF7 MX-100 | Human | Luminal | + | + | - | Metastatic Adenocarcinoma | Gifted by A.M Yu |
| 4T1 | Mouse | Luminal, basal | - | - | - | Metastatic Adenocarcinoma | ATCC #CRL-2539 |
| Met-1 | Mouse | Luminal | - | unk | unk | Metastatic Adenocarcinoma | Gifted by A. Borowsky |
| NDL | Mouse | Luminal | - | - | + | Invasive Ductal Carcinoma | Derived from primary tumor |
| UCDK9 MM3 | Dog | unk | unk | unk | unk | Melanoma | Gifted by R. Rebhun |
| UCDK9 OSA29 | Dog | unk | unk | unk | unk | Osteosarcoma | Gifted by R. Rebhun |
| D17 | Dog | unk | unk | unk | unk | Osteosarcoma | Gifted by R. Rebhun |

**Supplementary Table S1. Sources and characteristics of cancer cell lines employed in this study.** Cell lines from human, mouse and dog origins were previously derived from the indicated tumor tissue types and classified based on cell of origin (basal or luminal) and hormone receptor expression status (estrogen receptor (ER), progesterone receptor (PR), and epidermal growth factor receptor 2/human epidermal growth factor receptor 2 (HER2). Receptor overexpression is shown as positive (+), negative (-) or unknown (unk). Several cell lines were purchased from American Type Culture Collection (ATCC).

| Compound | Final Concentration | Solvent | Source |
|---|---|---|---|
| 10357 | 0-500 μM | DMSO | Gifted by F.A. Gorin |
| Amiloride | 0-500 μM | DMSO | #A7410, Sigma Aldrich |
| Benzamil | 0-500 μM | DMSO | #B2417, Sigma Aldrich |
| HMA | 0-100 μM | DMSO | #A9561, Sigma Aldrich |
| LLC1 | 0-100 μM | DMSO | Synthesized by CCCBSR |
| LLC2 | 0-100 μM | DMSO | Synthesized by CCCBSR |
| LLC3 | 0-100 μM | DMSO | Synthesized by CCCBSR |
| LLC4 | 0-100 μM | DMSO | Synthesized by CCCBSR |
| LLC5 | 0-100 μM | DMSO | Synthesized by CCCBSR |
| LLC7 | 0-100 μM | DMSO | Synthesized by CCCBSR |
| LLC8 | 0-100 μM | DMSO | Synthesized by CCCBSR |
| LLC9 | 0-100 μM | DMSO | Synthesized by CCCBSR |
| LLC10 | 0-500 μM | DMSO | Synthesized by CCCBSR |
| Docetaxel | 50nM – 10μM | DMSO | #S1148, Selleckchem |
| Doxorubicin | 75nM – 10μM | $H_2O$ | #BP25165, Fisher Scientific |
| (Z)-4-Hydroxytamoxifen | 250nM – 10μM | EtOH | #H7904, Sigma Aldrich |

**Supplementary Table S2. Characteristics of drugs employed in this study.** The final concentrations, solvents, and sources of all agents are summarized. Abbreviations used: dimethylsulfoxide (DMSO), ethanol (EtOH). LLC1 through LLC10 were synthesized, purified and characterized by the Combinatorial Chemistry and Chemical Biology Shared Resource (CCCBSR) of the UC Davis Comprehensive Cancer Center.

| Murine | HMA (μM) | LLC1 (μM) | Canine | HMA (μM) | LLC1 (μM) |
|---|---|---|---|---|---|
| Met-1 | 24 ± 4 | 7 ± 0.1 | UCDK9MM3 | 35 ± 9 | 11 ± 2 |
| NDL | 29 ± 6 | 8 ± 0.6 | UCDK9OSA29 | 25 ± 4 | 12 ± 2 |
| 4T1 | 24 ± 2 | 9 ± 1.6 | D17 | 42 ± 2 | 14 ± 1 |

**Supplementary Table S3. HMA and LLC1 are effective toward murine mammary tumor and canine cancer cell lines.** Several murine and canine cell lines were titrated with LLC1 or HMA for 24hr to determine viability via MTT assay. $IC_{50}$ values are shown in micromolar. Data represent three biological replicates.

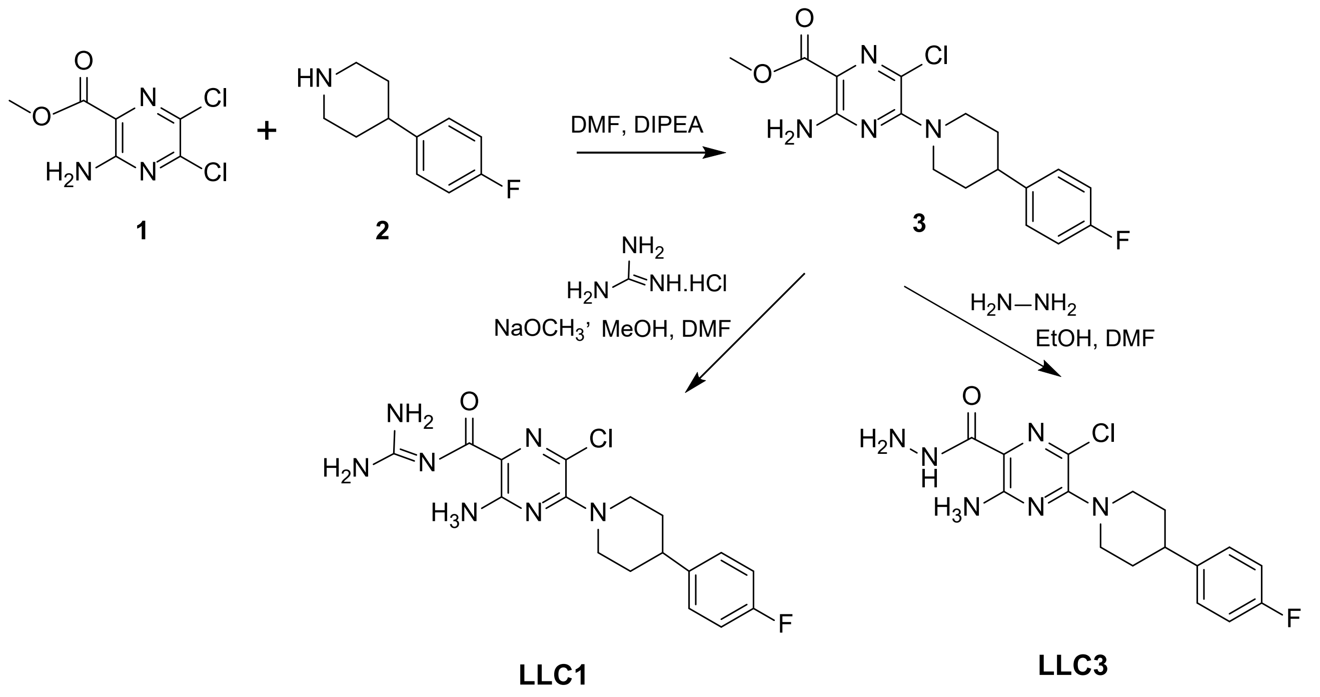


**Supplementary Figure S1.** Synthetic scheme for LLC1 and LLC3.

A

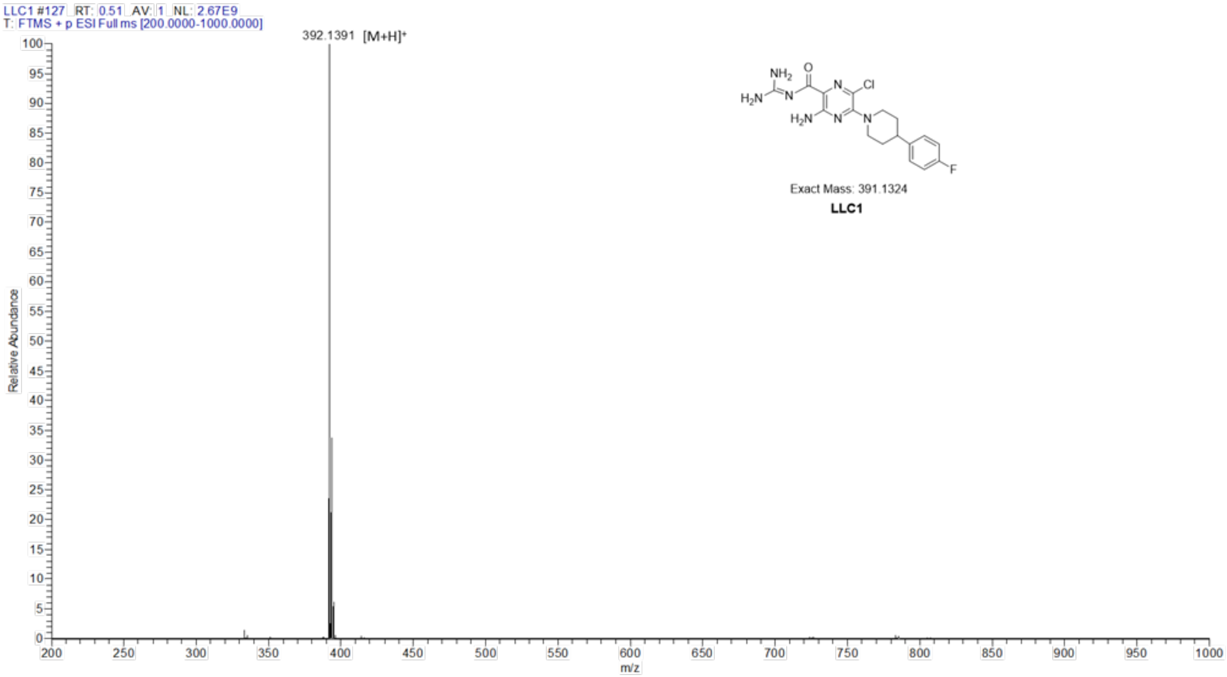


B

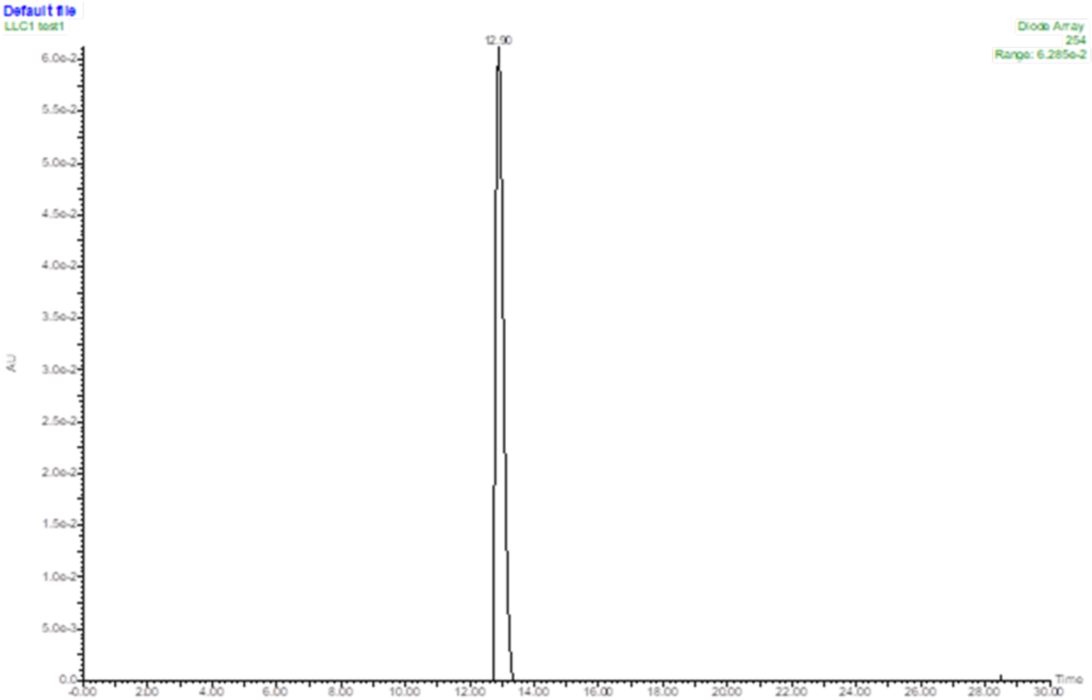


**Supplementary Figure S2. (A)** ESI-MS of LLC1. ESI-MS confirmed the structure of LLC1: calculated: m/z 392.1402, found: 392.1391 $[M+H]^+$. **(B)** HPLC of LLC1. HPLC chromatogram showed purity of LLC1 was > 98%**.**

**A**

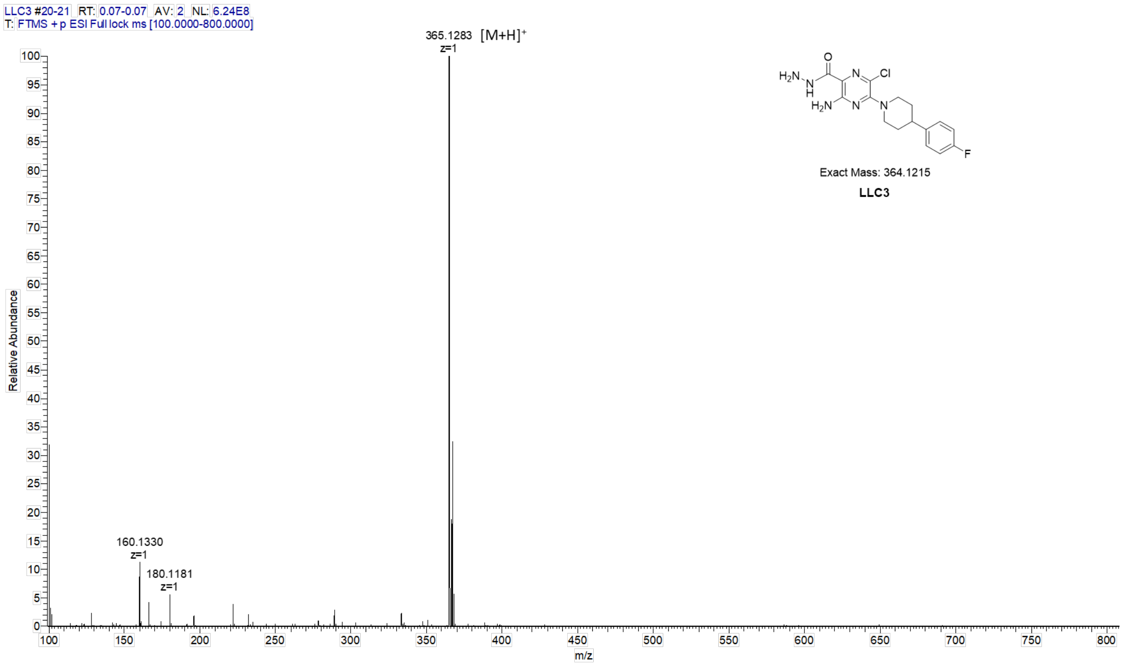


**B**

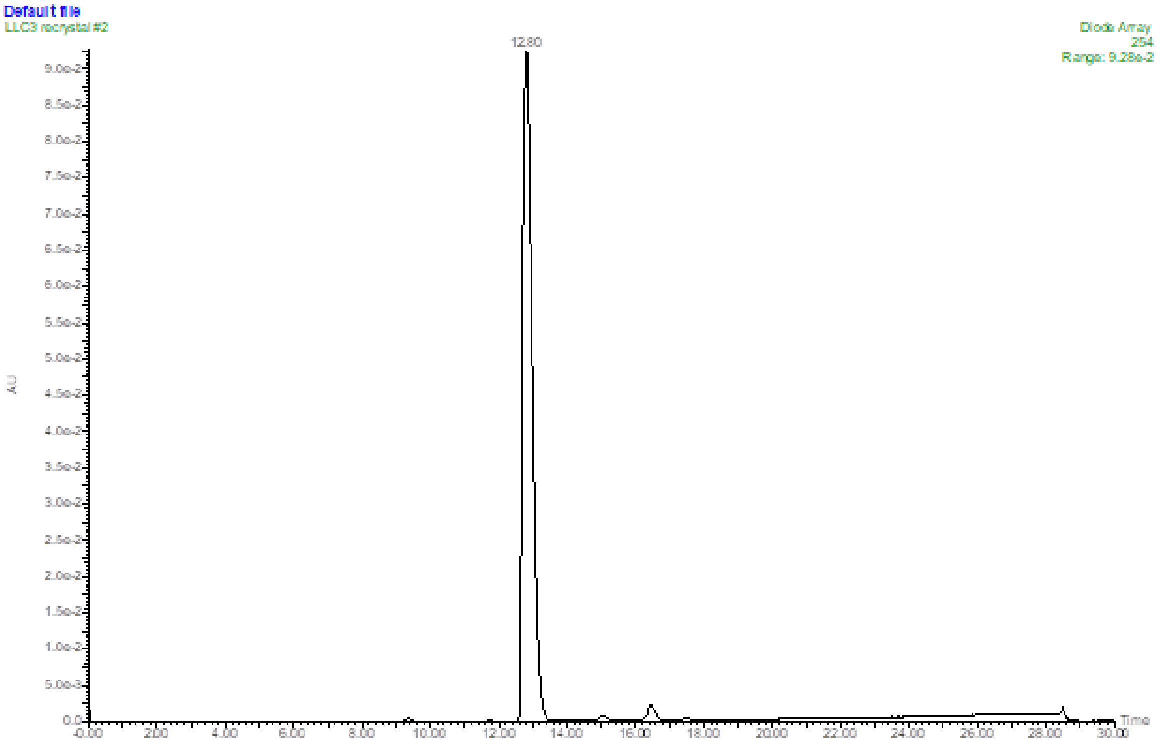


**Supplementary Figure S3. (A)** ESI-MS of LLC3. ESI-MS confirmed the structure of LLC3: calculated: m/z 365.1293, found: 392.1283 [M+H]$^+$. **(B)** HPLC of LLC3.

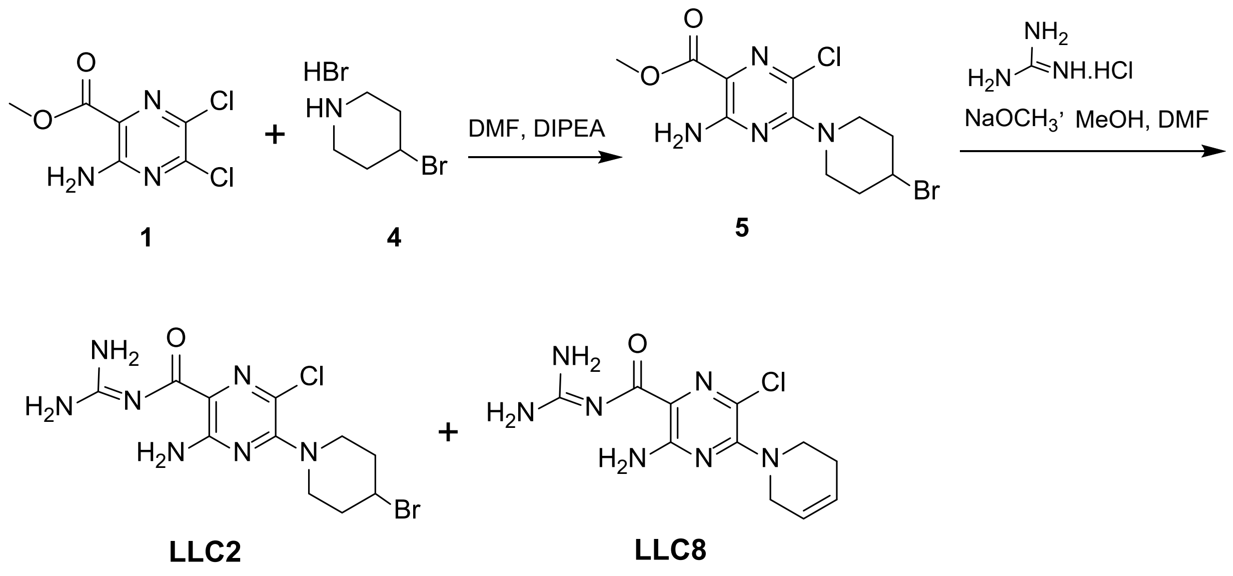


**Supplementary Figure S4.** Synthetic scheme for LLC2 and LLC8.

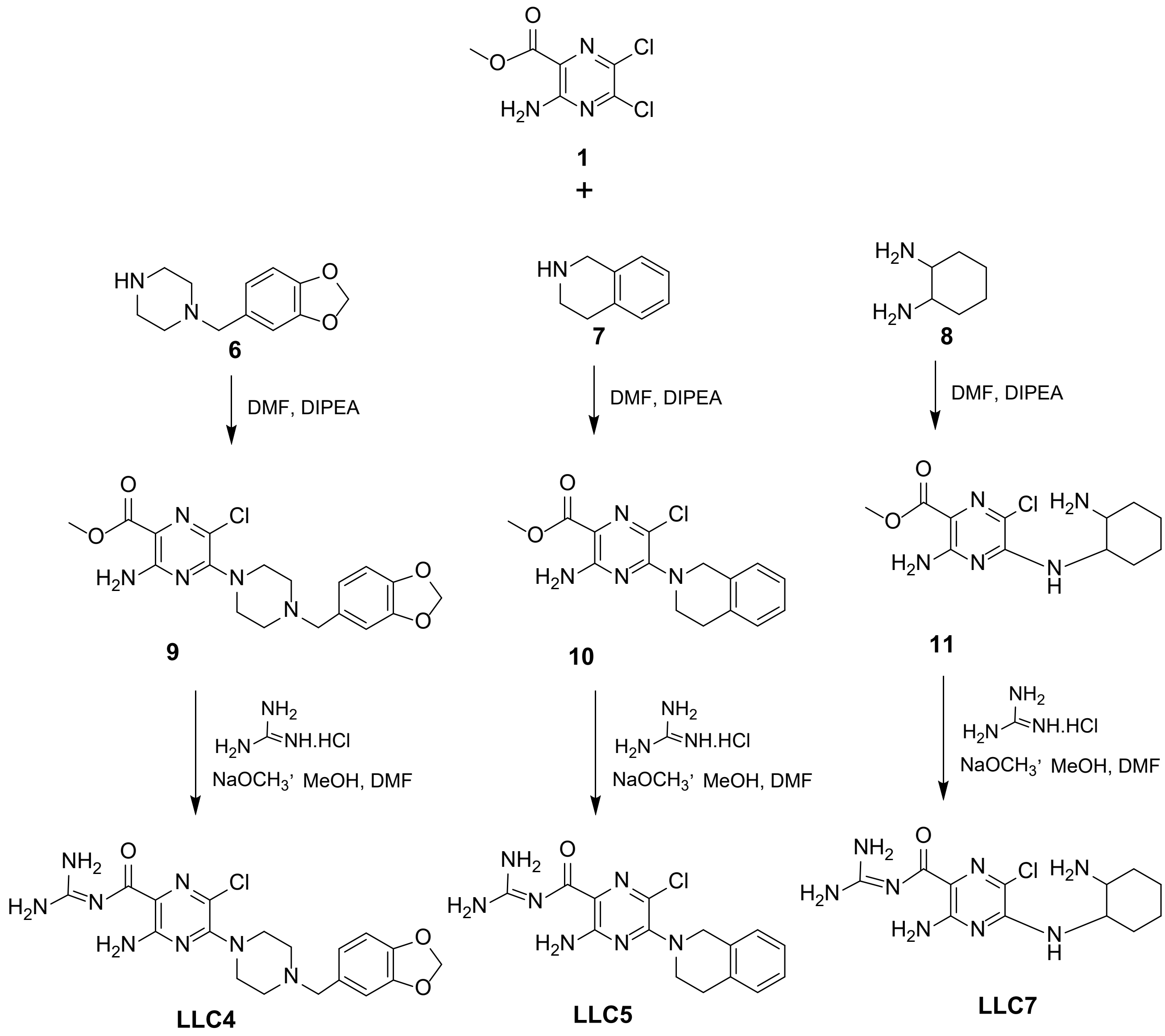


**Supplementary Figure S5.** Synthetic schemes for LLC4, LLC5 and LLC7.

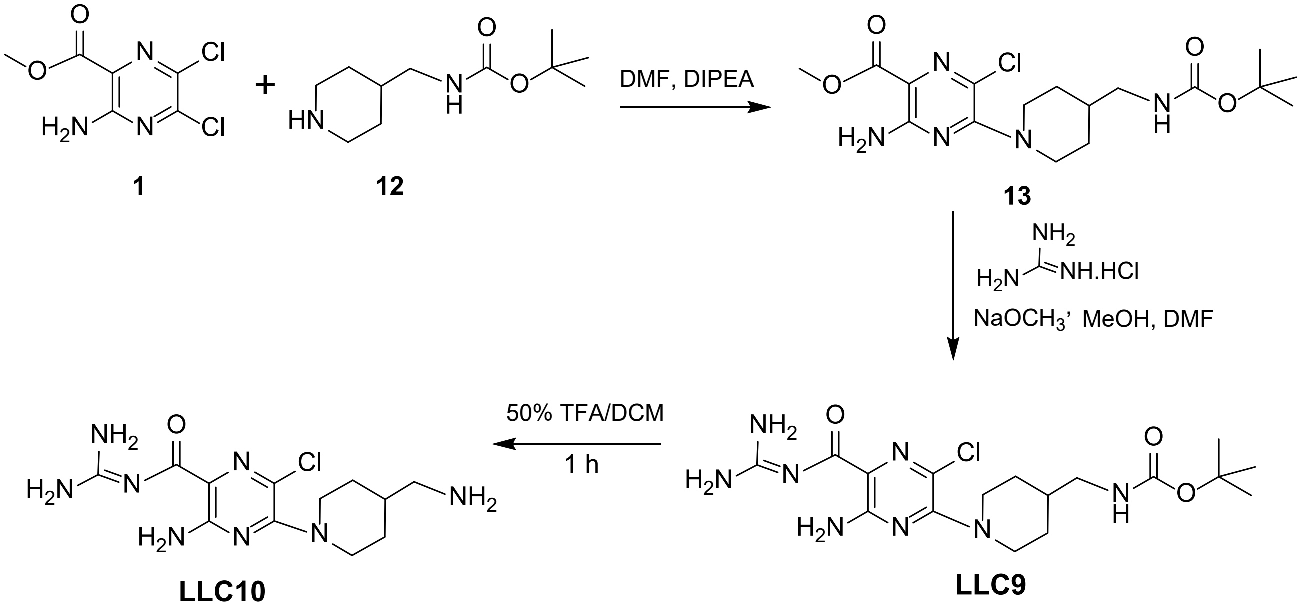


**Supplementary Figure S6.** Synthetic schemes for LLC9 and LLC10.

**A**

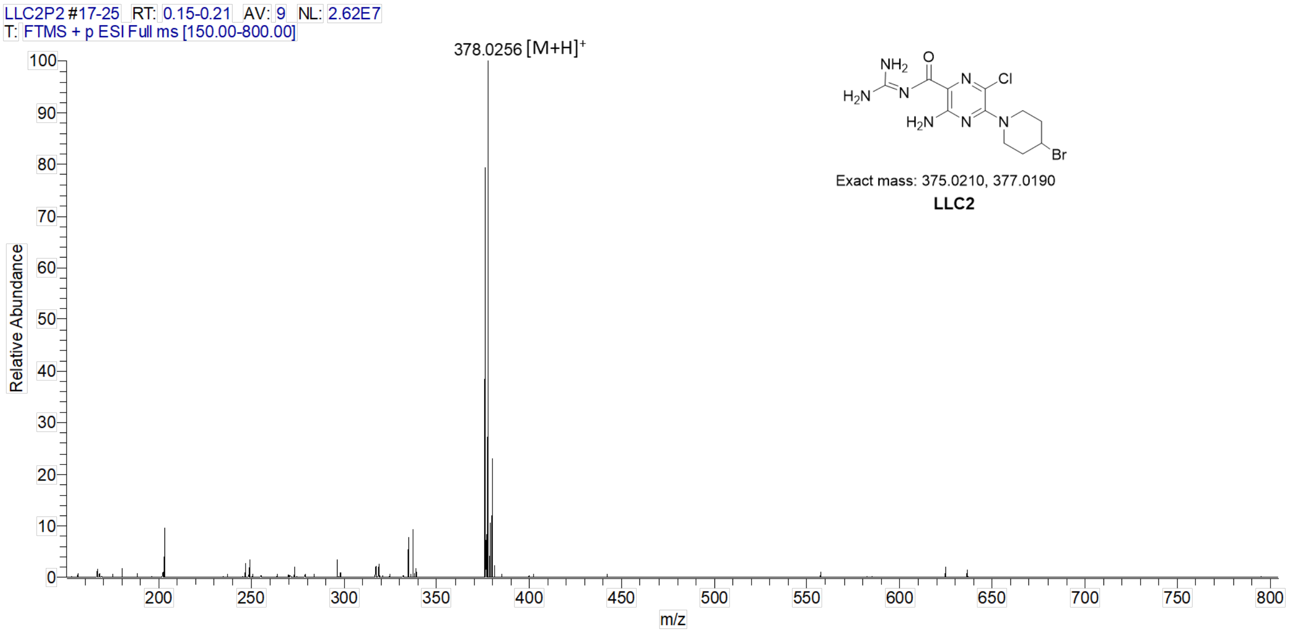


**B**

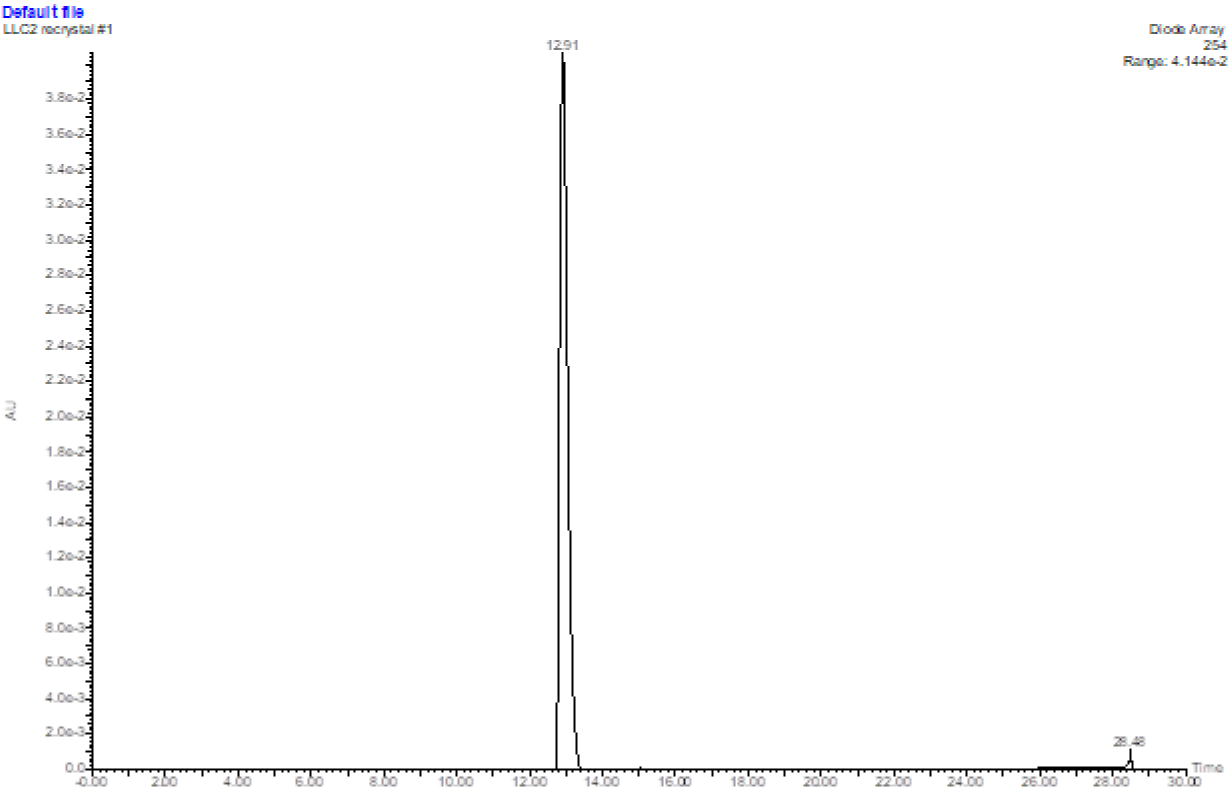


**Supplementary Figure S7. (A)** ESI-MS of LLC2. ESI-MS confirmed the structure of LLC2: calculated: m/z 376.0288, 378.0268, found: 376.0274, 378.0256 $[M+H]^+$. **(B)** HPLC of LLC2.

**A**

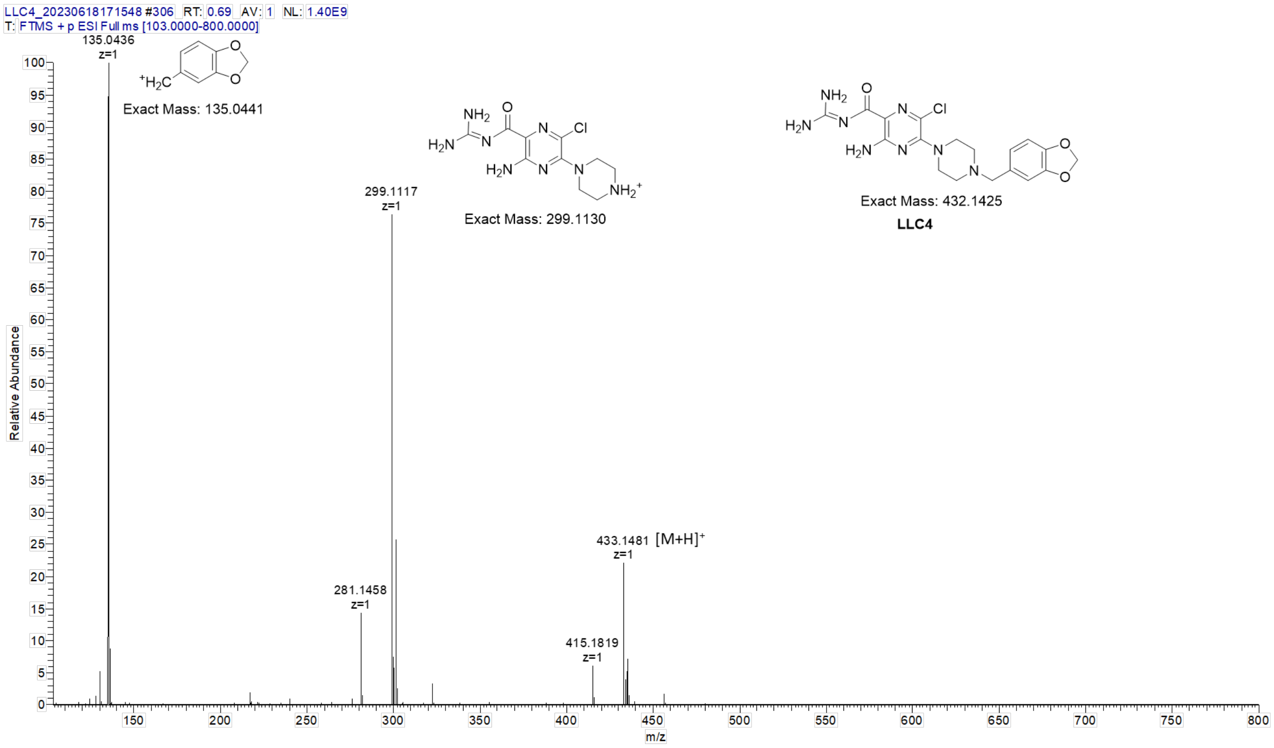


**B**

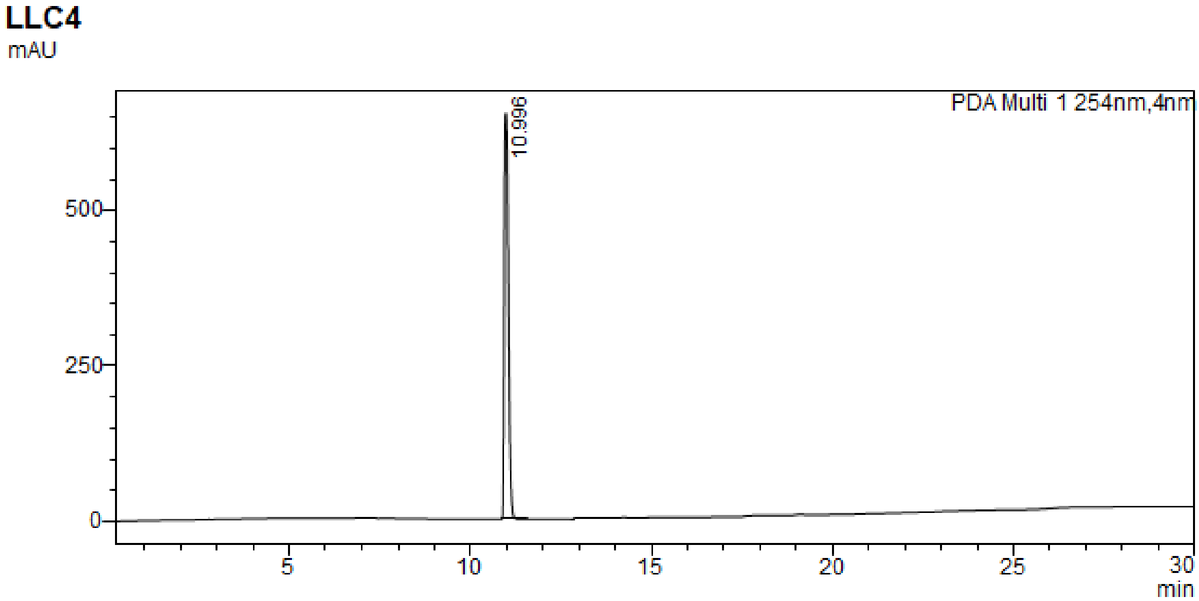


**Supplementary Figure S8. (A)** ESI-MS of LLC4. ESI-MS confirmed the structure of LLC4: calculated: m/z 433.1503, found: 433.1481 $[M+H]^+$, fragment peaks: calculated: m/z 135.0441, found: 135.0436; calculated: m/z 299.1130, found: 299.1117. **(B)** HPLC of LLC4.

**A**

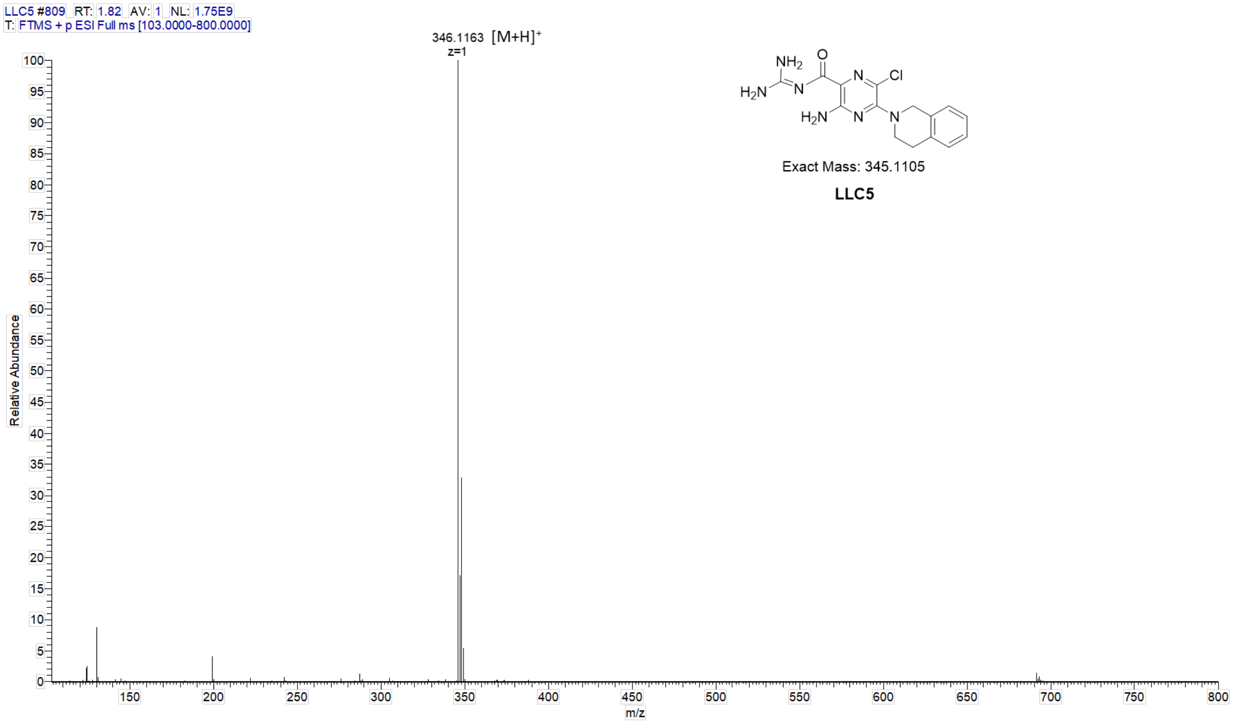


**B**

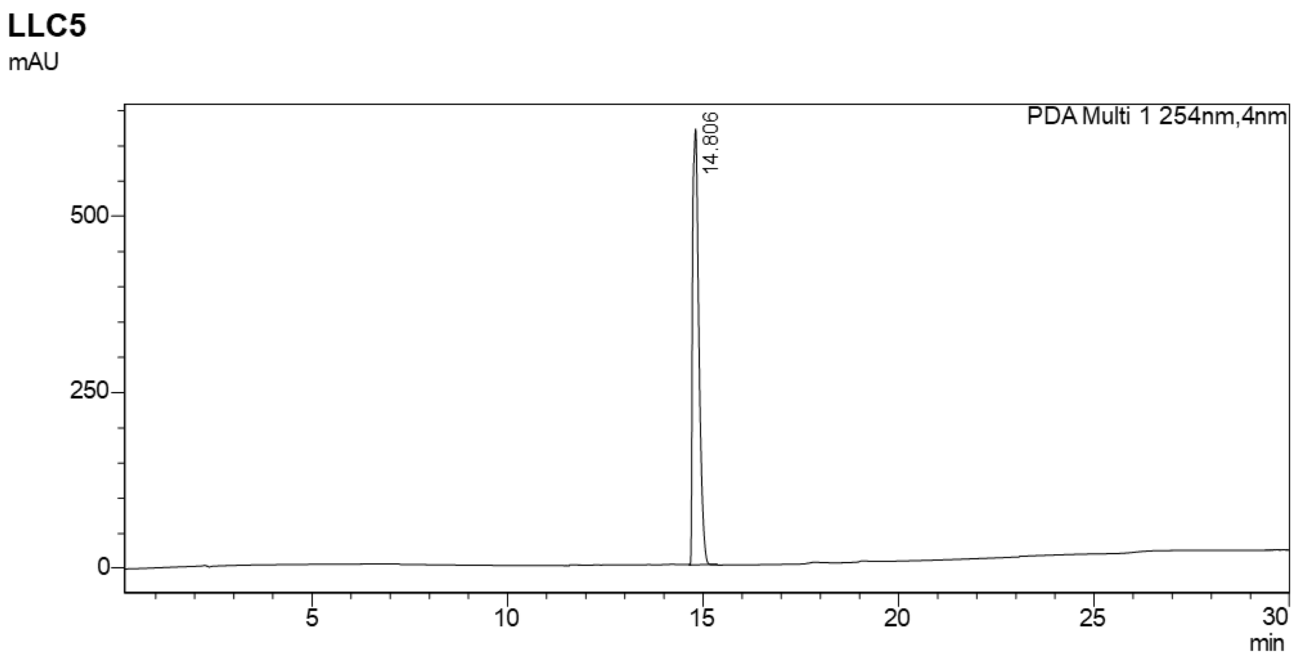


**Supplementary Figure S9. (A)** ESI-MS of LLC5. ESI-MS confirmed the structure of LLC5: calculated: m/z 346.1183, found: 346.1163 $[M+H]^+$. **(B)** HPLC of LLC5.

**A**

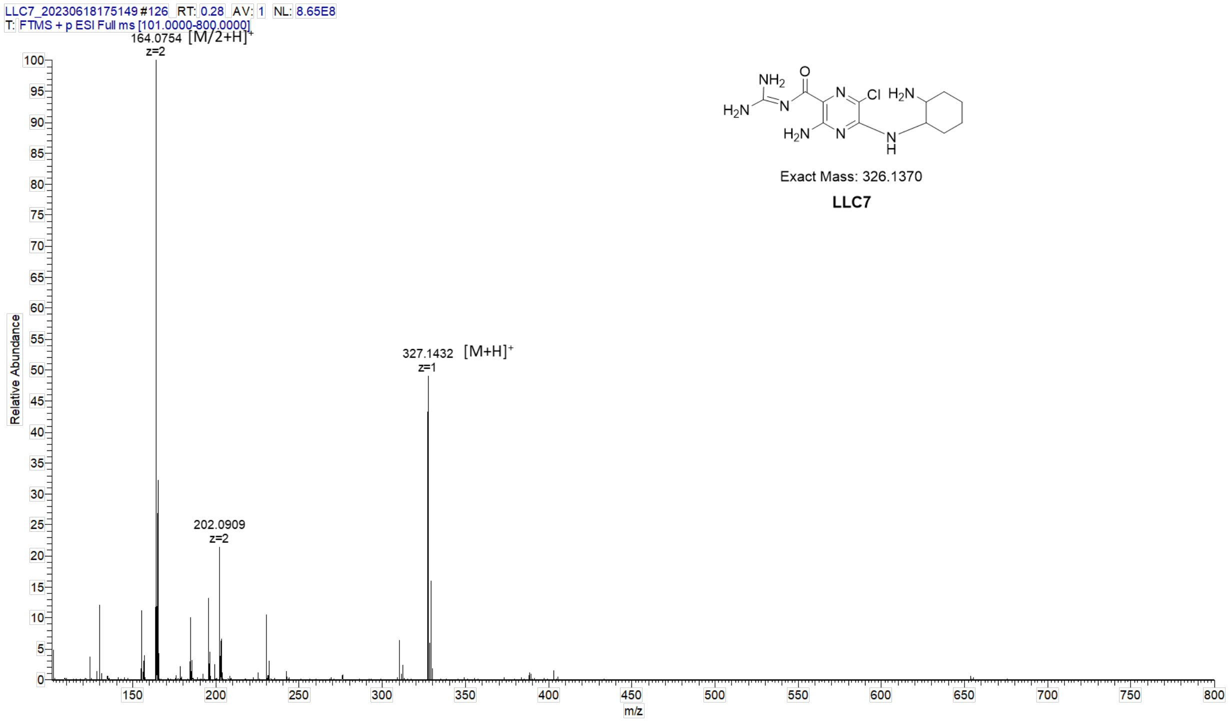


**B**

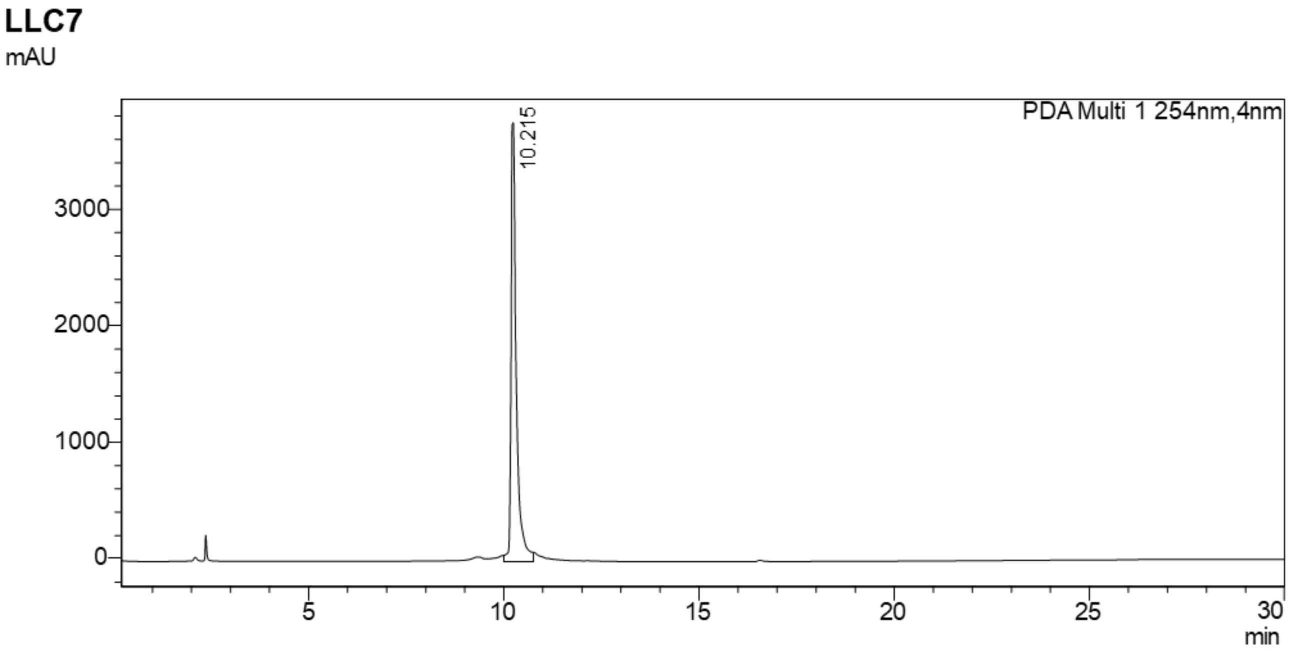


**Supplementary Figure S10. (A)** ESI-MS of LLC7. ESI-MS confirmed the structure of LLC7: calculated: m/z 327.1449, found: 327.1432 $[M+H]^+$, 164.0754 $[M/2+H]^+$. **(B)** HPLC of LLC7.

**A**

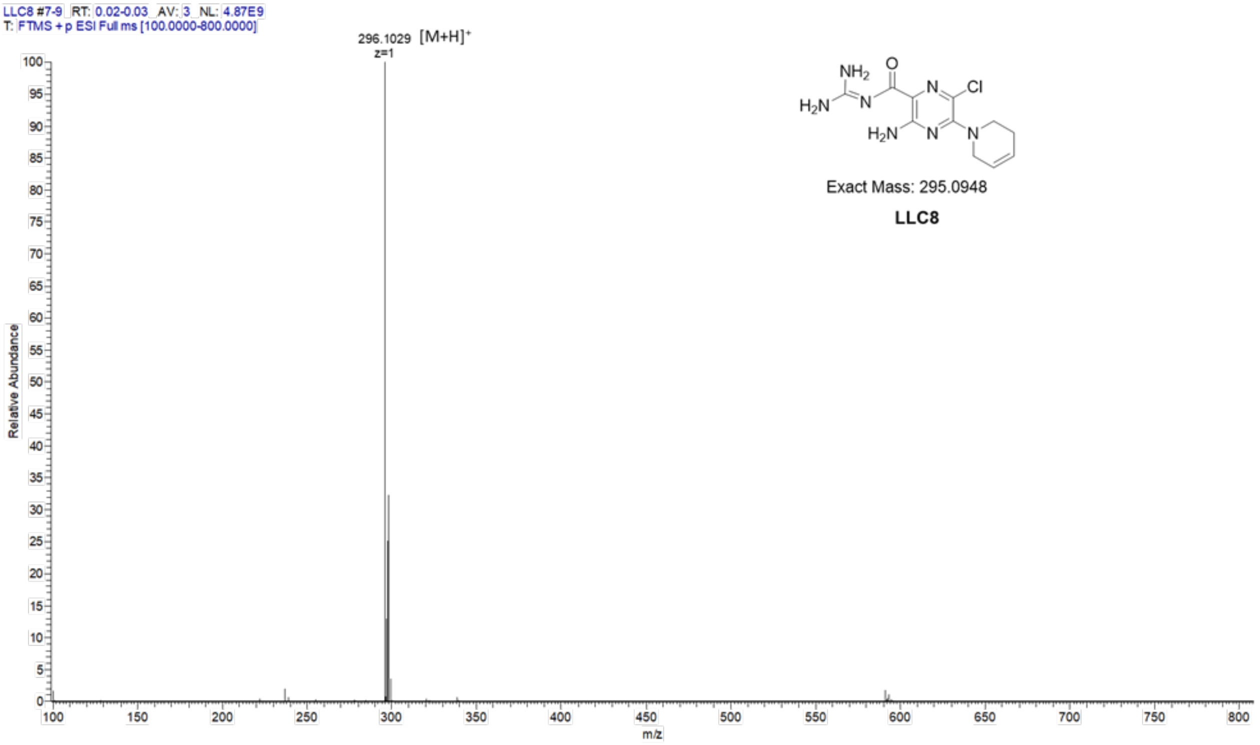


**B**

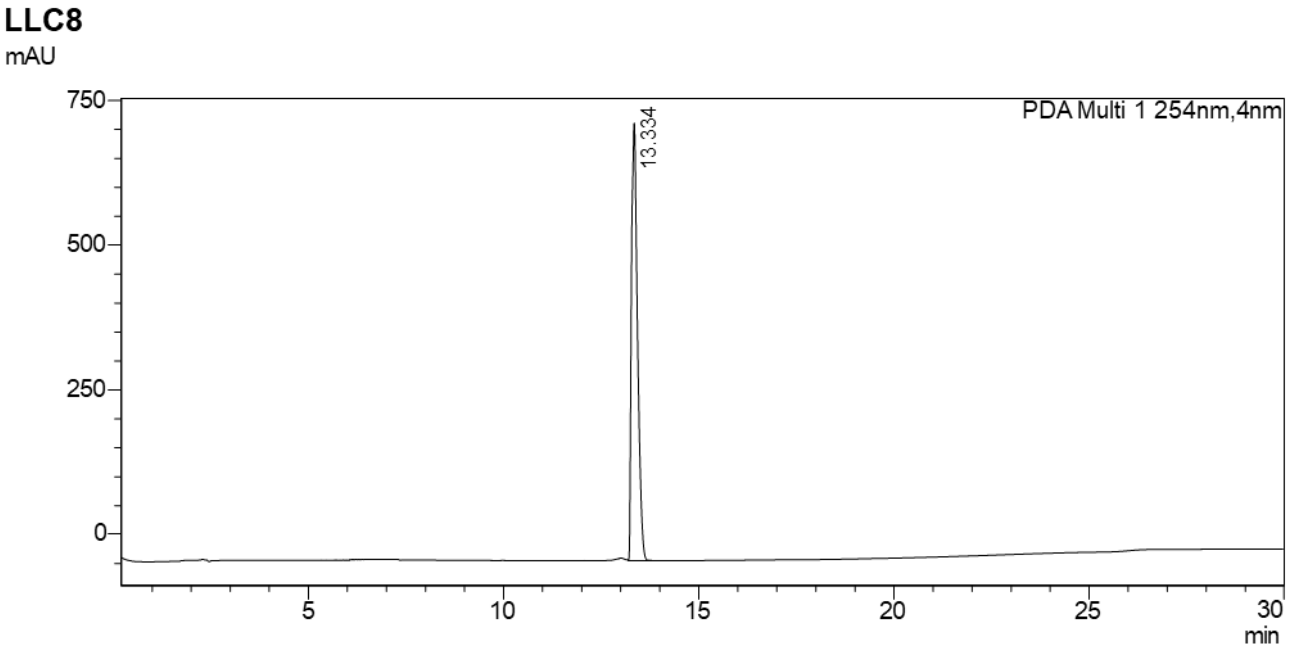


**Supplementary Figure S11. (A)** ESI-MS of LLC8. ESI-MS confirmed the structure of LLC8: calculated: m/z 296.1027, found: 296.1029 $[M+H]^+$. **(B)** HPLC of LLC8.

A

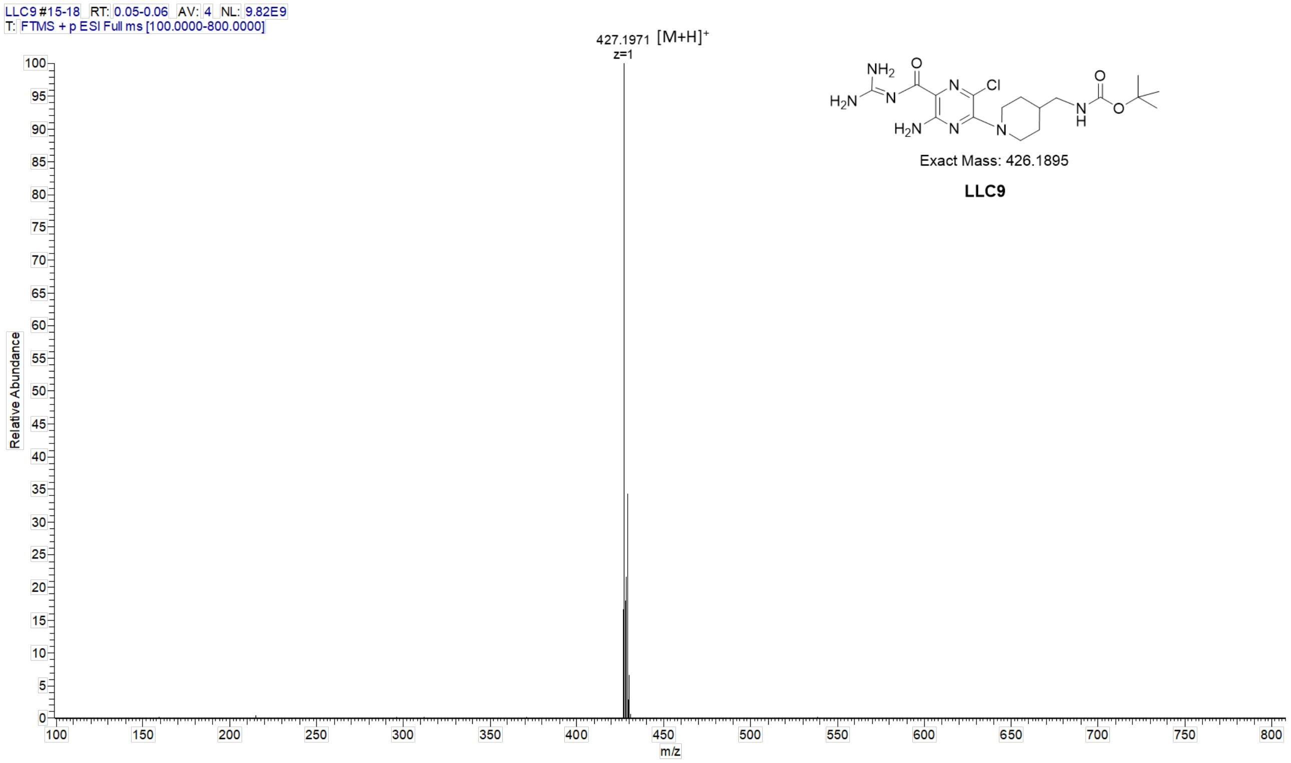


B

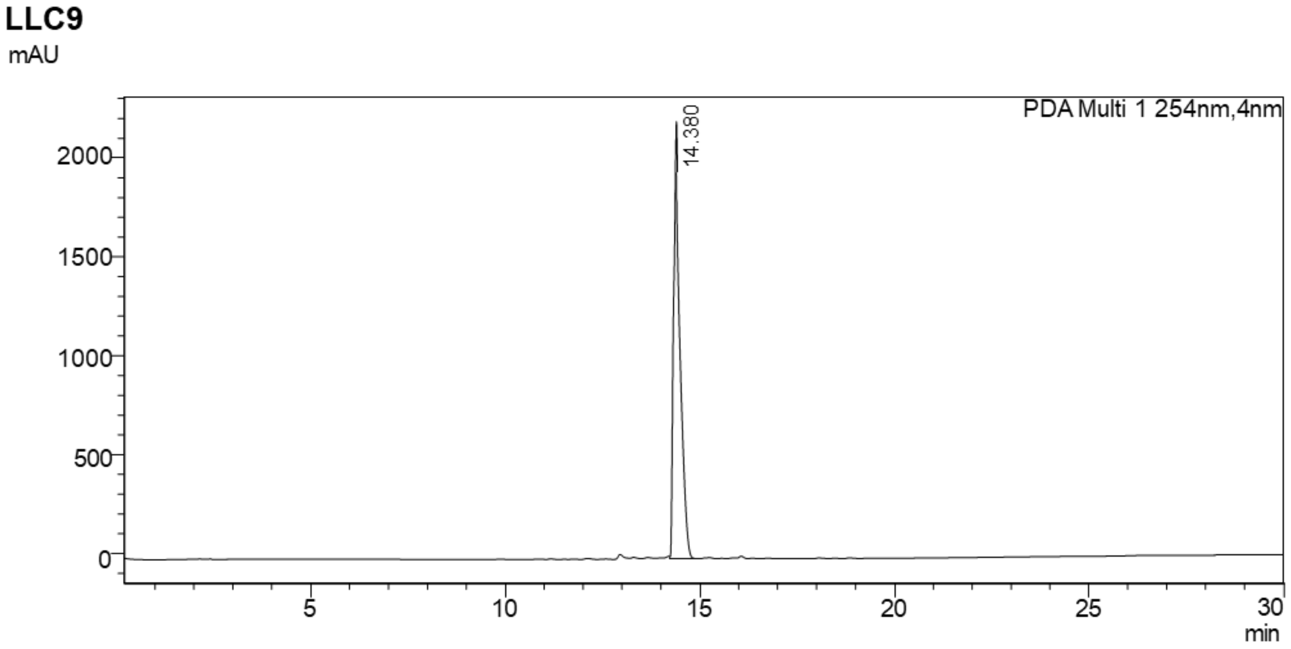


**Supplementary Figure S12. (A)** ESI-MS of LLC9. ESI-MS confirmed the structure of LLC9: calculated: m/z 427.1973, found: 427.1971 $[M+H]^+$. **(B)** HPLC of LLC9.

**A**

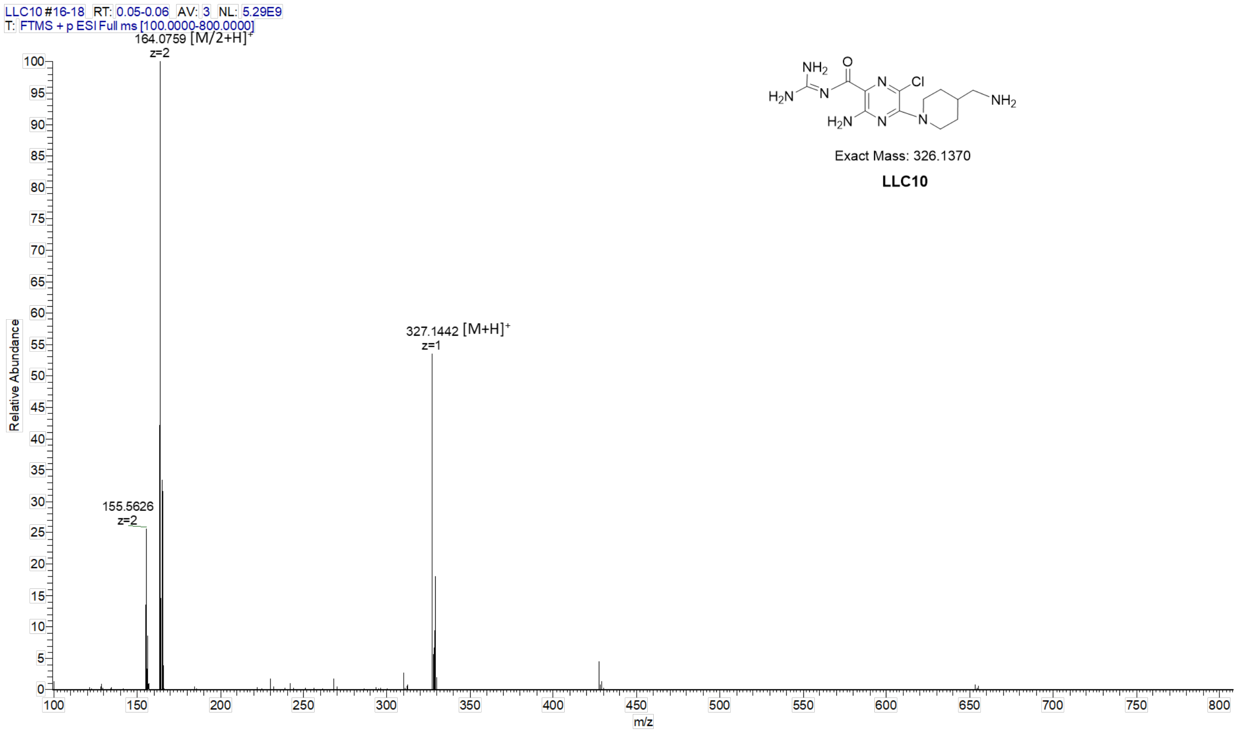


**B**

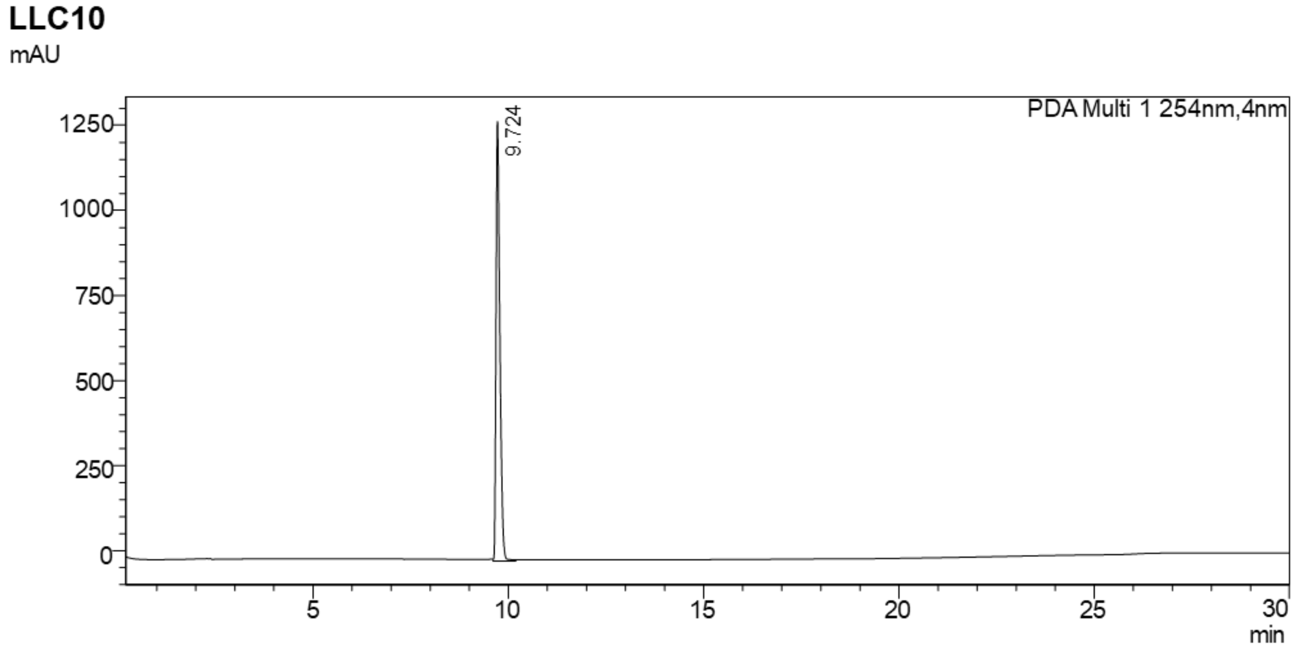


**Supplementary Figure S13. (A)** ESI-MS of LLC10. ESI-MS confirmed the structure of LLC10: calculated: m/z 327.1449, found: 327.1442 $[M+H]^+$. 164.0759 $[M/2+H]^+$. **(B)** HPLC of LLC10.

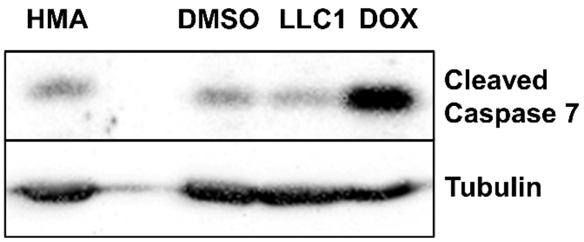


**Supplementary Figure S14. LLC1- and HMA do not provoke apoptotic cell death.** MDA-MB-231 cells were treated for 24hr as indicated with DMSO, 40 μM HMA, or 10 μM LLC1, or for 72hr with 10 μM doxorubicin as a positive control for drug-induced apoptosis. Lysates were immunoblotted with antibodies for cleaved caspase 7 (Cell Signaling, 9491) and tubulin (Sigma, T5168).

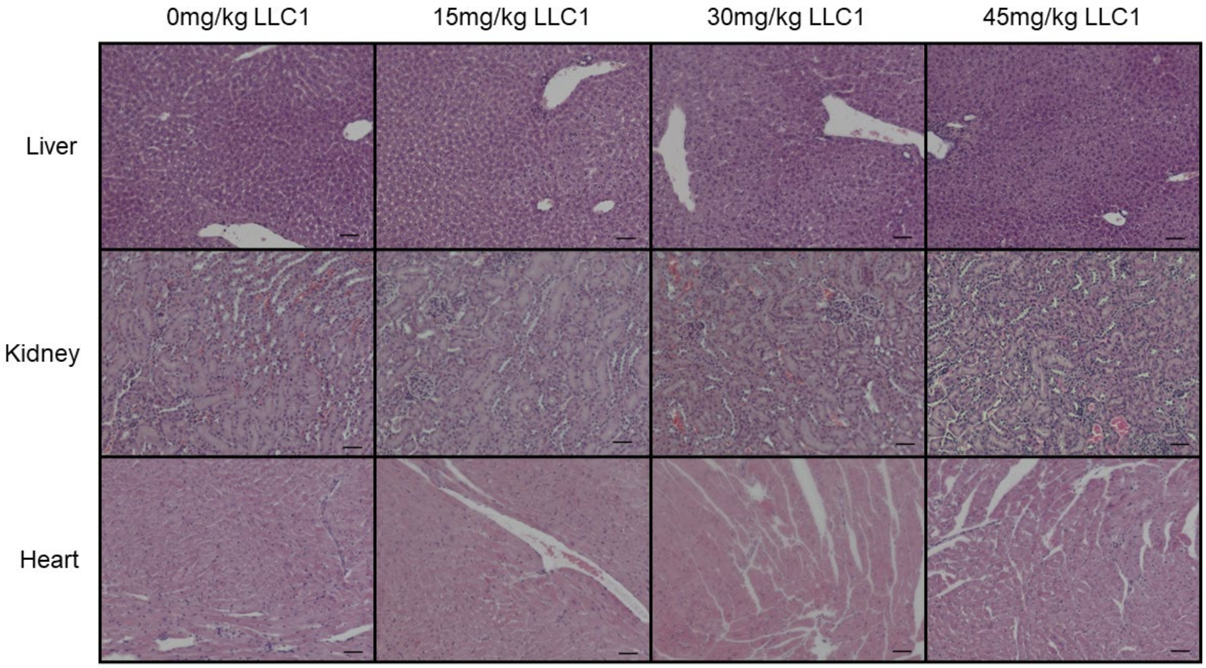


**Supplementary Figure S15. Histopathology of tissues after LLC1 treatment.** H&E staining of 5μm paraffin-embedded tissue sections of liver, kidney, and heart from the maximum tolerated dose experiment is depicted (scale bar = 50μm).

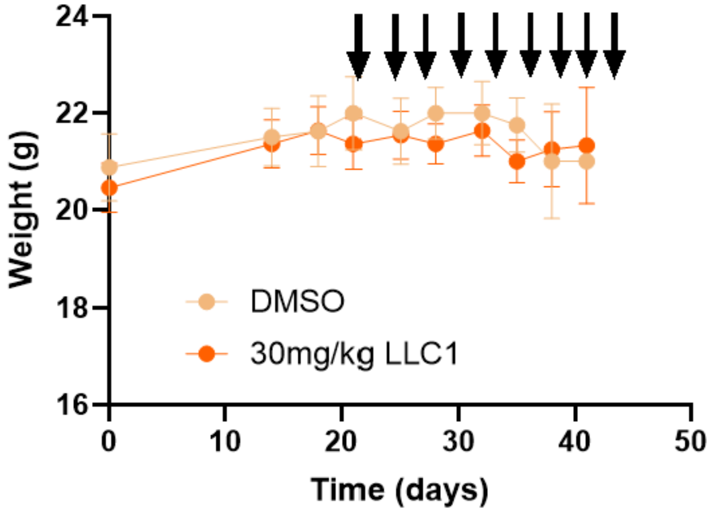


**Supplementary Figure S16. Body weights of mice throughout the therapeutic study.** Arrows depict treatment days.
